# Neuroinflammation driven by TLR7 activation in mice results in a global inflammatory response driving circuit-specific changes in neuronal gene expression

**DOI:** 10.1101/2025.04.25.650583

**Authors:** Kirstyn Gardner-Stephen, Robin J Carvajal-Quisilema, Lilya Andrianova, Rhona McGonigal, Deepika Sharma, Jennifer A Barrie, Megan Saathoff, John J Cole, Nigel B Jamieson, Michael T Craig, Jonathan T Cavanagh

## Abstract

Interactions between the brain and immune system play a key role in the aetiology of brain disorders, with inflammation emerging as a potential causal factor in subsets of major depressive disorder, particularly those resistant to treatment. The causal mechanisms through which immune activation can drive depressive symptoms remain elusive, limiting the ability to develop new targeted therapies. Using a mouse model of neuroinflammation, involving a TLR7/8 agonist, we found central and systemic inflammation alongside anhedonia-like behaviours, altered thalamostriatal signalling and infiltration of peripheral immune cells into the brain. Here, we sought to use combined whole-brain transcriptome and spatial transcriptomics approaches to determine whether Aldara-driven neuroinflammation resulted in consistent immune and neurobiological changes throughout the brain. We found evidence of strong immune activation throughout the brain, with astrocytes displaying a strong inflammatory profile that was relatively uniform throughout. However, we found that this global inflammatory signal led to regionally-specific changes in gene expression, particularly reduced expression of genes associated with synaptic function in brain areas underlying mood and anxiety, such as ventral striatum and amygdala. Our data suggest potential mechanisms through which astrocytes regulate neuronal function in response to inflammation.

## Introduction

Brain-immune interactions are implicated in the aetiology of many neuropsychiatric conditions including major depressive disorder (MDD)^1, 2^. Anhedonia, the loss of motivation and ability to feel pleasure, is a core symptom experienced by many individuals with mood disorders^3^. Human brain imaging studies reveal that anhedonia is associated with changes in the brain’s reward circuitry, particularly interactions between midline thalamus and ventral striatum^4, 5^. We have also recently reported that, in both individuals with inflammatory disease and in a preclinical mouse model of neuroinflammation, altered connectivity between thalamus and striatum is associated with depression in humans and anhedonic behaviour in mice^6^.

While accumulating evidence shows that neuroinflammation can induce behavioural changes by altering the function of reward circuitry in the brain, the specific causal mechanisms through which neuroinflammation can alter neural function remain poorly understood. Neuroinflammation is associated with immune cell infiltration into the brain^7^, and we found that treatment with a topical TLR7/8 agonist, Aldara, led to ingress of cytokine-producing immune cells into the brain, including CD4+ and CD8+ T cells and monocytes^6^. In the neuroinflammatory environment microglia become reactive, undergoing morphological changes and produce proinflammatory cytokines^8^. This cytokine production by microglia in turn induces astrocytes to adopt a reactive phenotype that can impair neural circuit function^9^. Indeed, astrocytes have been implicated in modulating neuroinflammatory environments by mediating crosstalk between CNS and infiltrating immune populations^10, 11^.

Astrocytes play a key role in regulating many aspects of neural circuit function, including neurotransmitter cycling, ion balance and providing trophic support to neurons, as well as neurovascular coupling and maintaining the blood-brain-barrier^12^. Alterations in astrocytic function by neuroinflammation can therefore have a profound impact on neural circuit function^11^. Using the preclinical rodent model of Aldara-driven neuroinflammation, we found strong evidence of both microglial and astrocytic reactivity with proinflammatory cytokine production, in addition to anhedonia-like behaviours and disrupted glutamatergic signalling from midline thalamus to ventral striatum^6^. Previous work with this model revealed that the TLR7-activated inflammation results in an interferon (IFN)-mediated inflammatory cascade that induces pro-inflammatory dermal and systemic responses in addition to neuroinflammation^13, 14, 15^.

Data from this model included evidence of widespread neuroinflammation and suggested that changes in neural circuit function were limited to the reward circuitry^6^ The aim of the present study was to determine whether the consequences of Aldara-driven neuroinflammation are regionally-specific. We approached this using an unbiased spatial transcriptomics approach. To characterise this model at whole tissue, cellular and spatial molecular level, we integrated whole-brain bulk RNA sequencing with Nanostring CosMx™ spatial molecular imager (SMI) platform^16^. This spatial transcriptomics technique allows for single cell, subcellular and spatially resolved transcriptional findings in respect to the 1000-plex RNA Mouse Neuroscience panel across entire mouse brain sections. This transcriptional exploration of the TLR7-mediated neuroinflammatory model provided both global and spatially resolved data on the astrocytic contribution to neuroinflammation and changes to neurotransmission-related gene expression.

## Methods

### Animals

Female C57BL/6J mice (8 weeks old) were obtained from Charles River Laboratories (UK) and group-housed in ventilated cages. Animals underwent acclimatisation for one week prior to experimental use. Animals were maintained in a 12 h light/dark cycle in controlled temperature and humidity with *ad libitum* access to food and water. All experiments were carried out in accordance with the UK Animal (Scientific Procedures) Act 1986 and were subject to local ethical approval by the University of Glasgow Animal Welfare and Ethical Review Board.

### Aldara Imiquimod (IMQ) model

A 3cm^2^ area was shaved on the dorsal back region of adult mice as per previous studies ^13^. Mice then underwent three consecutive days of treatment with either 62.5mg of Aldara^TM^ cream (5% IMQ: Meda AB, Sweden) or Control cream (Boots Aqueous Cream B.P., Boots Pharmacy, UK) applied to the shaved area. Psoriasis-like dermal inflammation was assessed daily using a scoring matrix for inflammation, erythema and scaling caused by topical Aldara cream application. Bulk RNA sequencing experiments used 5 mice per group, while CosMx™ spatial transcriptomics experiments used 4 mice per group.

### Tissue harvesting and sectioning

Mice were terminally anaesthetised via intraperitoneal injection of pentobarbital (Euthatal, Dechra, 10 μl/g of 200 mg/ml solution) and transcardially perfused with 20ml of ice-cold PBS (Merck, UK) to remove residual peripheral blood from the brain vasculature. Brain tissue was then harvested as per the requirements of the proceeding experiments all under RNAase-free conditions.

### RNA extraction and bulk RNA sequencing

Brains were hemisected in RNAse-free conditions and the hemispheres were moved to Qiazol Lysis Reagent (Qiagen, UK) and homogenized using a TissueLyser LT (Qiagen, UK) for three minutes with a five mm stainless steel ball (Qiagen, Cat. No. 69989, UK). RNA was isolated using the Qiagen RNeasy Lipid Tissue Extraction Kit (Qiagen, Cat. No. 74804, UK) with RNeasy on-column treatment as per manufacturer’s instructions. RNA quality was assessed using an Agilent 2100 Bioanalyzer (Agilent Technologies) −80°C until being sent for bulk RNAseq processing. A minimum of 7 μg per sample was sent to Novogene. Library preparation and bulk RNA sequencing was completed by Novogene, with a read length of 2×150 bp and an average of 53 million reads per sample. The data was aligned to the mouse genome (ensembl_mus_musculus_grcm38_p6_gca_000001635_8) using HiSat2, with > 90% uniquely aligned concordant pairs per sample. Raw read counts were normalised and differential expression calculated using DESeq2 ^17^ using pairwise models with no additional covariates. The normalised data were explored using the Searchlight analysis platform, with the pipeline described in detail elsewhere^18^. The significance threshold for differential genes was adjusted p < 0.05 and absolute log_2_fold change > 0.5. All other parameters were left at default.

Overrepresentation analysis (ORA) and visualisation was carried out using G:Profiler and the G:Profiler2 package^19^. For the ORA analysis in G:Profiler, we ordered the DE gene lists by most significant for upregulated and downregulated genes and run ORA using the g:GOSt function with this ordered input against the *Mus musculus* database. The significance threshold was set to adjusted p < 0.05 and only annotated genes were considered.

### FFPE tissue preparation and CosMx™ Mouse Neuroscience Panel

Mice were perfused with 20ml of ice-cold PBS followed by 20ml of ice-cold 4% paraformaldehyde (PFA) made in PBS (Sigma-Aldrich, UK), before brains were removed overnight into 4% PFA at 4°C. Following tissue dehydration via an ethanol concentration gradient, brains were paraffin embedded using a Thermo Scientific Shandon Citadel 1000 tissue processor (Thermo Fisher Scientific, UK) prior to paraffin embedding. Two mice from each experimental group were set in one paraffin block. For each biological replicate, we took a single coronal section in two anatomical planes: an anterior tissue plane (Bregma range = +1.1 to +0.78 mm) and posterior tissue plane (Bregma range = -1.0 to -2.0 mm).

Sections were mounted onto Leica Biosystems BOND Plus slides according to the CosMx imaging scan area boundaries. Tissue was then processed, stained and imaged using CosMx Mouse Neuroscience 1000-plex RNA Panel, as previously described^16^. Fields of view (FOVs) of 0.51mm^2^ were selected in a grid-like pattern to capture the entire tissue section on the CosMx SMI. A total of 1348 FOVs were selected across the 16 tissue samples.

### Data processing & quality control

A total of 16 coronal hemisections across 4 slides were analysed using CosMx SMI. Following image processing and decoding steps that were performed in Atomx, all slides and corresponding FOVs were exported as single-cell RNA-sequencing (scRNA-seq) data objects for analysis with Seurat toolkit, hereafter referred as Seurat objects^20^. All Seurat objects were exported with polygon coordinates and transcript coordinates, however the latter were excluded from downstream analysis and used only to visualise transcript localisation within segmentation polygons. All FOVs belonging to a single brain sample tissue were annotated with the same sample name, which was later used to generate individual objects using the subset function (Seurat v5.2.0), resulting in 16 Seurat objects derived from the original 4 slide-level Seurat objects. All of the generated objects were updated to Seurat version 5.2.0 using the UpdateSeuratObject function (Seurat v5.2.0), then merged and processed as one Seurat Object. Quality control and filtering were performed on the merged object to remove cells that express less than 20 transcripts, as per NanoString manufacturer recommendations. All the data processing and quality control steps were carried out using R programming Language (v4.4.1) ^21^ and RStudio (v2024.12.1+563) ^22^.

### Clustering and cell type identification

The merged Seurat object was log normalised using NormalizeData function (Seurat v5.2.0) and scaled using ScaleData function (Seurat v5.2.0). Principal component analysis (PCA) was performed using the RunPCA function (Seurat v5.2.0), with all genes from the 1000-plex panel considered as variable features across all samples. Integration was performed to align shared cell populations across samples and reduce technical variation. Batch correction was applied using Harmony^23^, implemented via the RunHarmony function (SeuratWrappers v0.3.0), with sample name and condition specified as batch variables (using the group.by.vars argument) and the diversity clustering penalty parameter (theta argument) set to 1 for both variables to balance correction strength with the preservation of biologically meaningful variation. Cell embeddings from the first 10 principal components were used to construct a shared nearest neighbour graph via the FindNeighbors function (Seurat v5.2.0), followed by community detection with the FindClusters function (Seurat v5.2.0) using the resolution argument set to 0.7. The harmony-corrected PCA embeddings were used for non-linear dimensionality reduction with Uniform Manifold Approximation and Projection via the RunUMAP function (Seurat v5.2.0). Differential gene expression analysis was performed using the FindAllMarkers function (Seurat v5.2.0) with Wilcoxon Rank Sum Test (test.use argument = “wilcox”) to identify cluster-defining marker genes. Cell type annotation was guided by the expression of canonical markers (Table 1) and spatial localisation patterns of clusters using ImageDimPlot function (Seurat v5.2.0).

**Table 1.** Marker genes used for cell type identification.

| **Cell type** | **Clusters** | **Cell identifier genes** |
| --- | --- | --- |
| **Glutamatergic Neurons** | 2, 6, 15 | *Slc17a7, Camk2a* |
| **Medium Spiny Neurons** | 8,13 | *Ppp1r1b, Adora2a, Arpp21,* |
| **GABAergic Neurons** | 3 | *Gad1, Gad2, Slc32a1* |
| **Oligodendrocytes** | 5,9 | *Mog, Mag, Plp1, Mbp* |
| **Astrocytes** | 7, 11, 12 | *Gja1, Slc1a3, Aqp4* |
| **Neurovascular** | 4 | *Plp1, Apod* |
| **Endothelial cells** | 14 | *Cldn5, Pecam1, Flt1* |
| **Ependymal** | 16 | *Foxj1, Ttr, Htcr2* |
| **Microglia/Immune** | 10, 17 | *Ptprc, Cx3cr1* |
| **Unclassified Neurons** | 0, 1 | - |

### Pseudo-bulk cell agnostic region of interest analysis

Brain images were exported from the integrated Seurat object as PNG files using the ImageDimPlot function (Seurat v5.2.0), with the boundaries argument set to “segmentation” and coloured by identified cell types (using the *group.by* argument). Segmentation boundaries were preferred over centroids as they better represented cell dimensions and avoided overlapping representations of adjacent cells. This approach was selected due to the spatial overlap observed between certain cell types and anatomical brain regions. Not all regions could be mapped onto the brain images due to limitations in anatomical landmark resolution.

Each brain slice image was exported at high resolution (3400 × 4250 pixels). Regions of interest (ROI) were annotated using a method that was later turned into the SubsetMask tool (code available on Github). The SubsetMask tool was developed to enable manual annotation of regions of interest on exported brain images. Black masks were overlaid onto the PNG images using image editing software, with one region defined per layer. The pixel coordinates corresponding to these masks were extracted and mapped back to the spatial coordinates of cells in the Seurat object, allowing annotation of specific regions directly within the dataset.

Anatomical reference was provided by the Allen Mouse Brain Atlas (Figure 4A). Regions of interest (ROI) were selected based on their established associations with inflammation-induced behavioural changes and included the anterior cingulate cortex (ACC), ventral striatum (including nucleus accumbens), thalamus, amygdala, and, as a control region, the motor cortex (including primary and secondary motor areas where applicable).

ROI mask contour coordinates were extracted using a custom script implemented with the OpenCV Python package (v4.11.0.86) ^24^. The extracted coordinates were scaled to the physical dimensions of each brain slice based on the minimum and maximum *x, y* values. Mask coordinates were converted into matrices and used to generate *Polygon* objects, which were subsequently wrapped into *SpatialPolygons* objects using the sp R package ^25^. Similarly, the *x, y* coordinates of each brain slice were transformed into matrices and converted into *SpatialPoints* objects. Cells whose coordinates overlapped with the corresponding *SpatialPolygons* were annotated in the metadata with their respective regions. Not all cells were labelled, as some regions could not be reliably identified, and padding was intentionally left between adjacent regions during ROI drawing to prevent misclassification of boundary cells.

### Transcriptomic analysis

Pseudo-bulking of single-cell and region data was performed using the AggregateData function (Seurat v5.2.0), grouping by cell type/region and sample name. Differential expression analysis was carried out using the DESeq2 R package^17^, with *condition* specified as the design variable. As anterior and posterior samples originated from the same mouse, analyses were performed separately for these regions to avoid over-sampling in the statistical comparisons. Transcriptomic analyses, including data visualization and pathway over-representation analysis, were conducted using the Searchlight2 pipeline^18^ incorporating STRING v11.5 ^26^ for functional enrichment. Analyses were performed at both the pseudo-bulk single-cell level and the regional level. The background organism for enrichment analyses was *Mus musculus*, using the Ensembl reference genome GRCm39 ^27^.

## Code Availability

The code is available at https://github.com/RobinCarvajal/cosmx_cavanagh and https://github.com/RobinCarvajal/SubsetMask

## Results

### Upregulation of immune-related genes in response to TLR7-induced neuroinflammation

Previously, we described neural transcriptional changes within the Aldara model by investigating selected inflammatory targets with qPCR^14 28^. We initially used a whole-brain RNAseq transcriptome approach to assay global changes in gene expression in response to Aldara treatment. We employed a principal component analysis (PCA), which revealed that the Aldara and control-treated mice clustered separately, indicating substantial differences in gene expression between the two groups (Figure 1A). Differential expression analysis using DESeq2^17^ revealed that there were 2198 significantly upregulated and 585 significantly downregulated genes in the brains of Aldara-treated mice, relative to control-treated mice (Figure 1B). To control over-representation of very highly expressed genes, all gene expression values were scaled on a gene-by-gene basis using the Z-score transformation, prior to the PCA being run. The significance threshold was set at p<0.05 (adjusted for FDR) and a minimum log fold change of 0.5.

**Figure 1:**
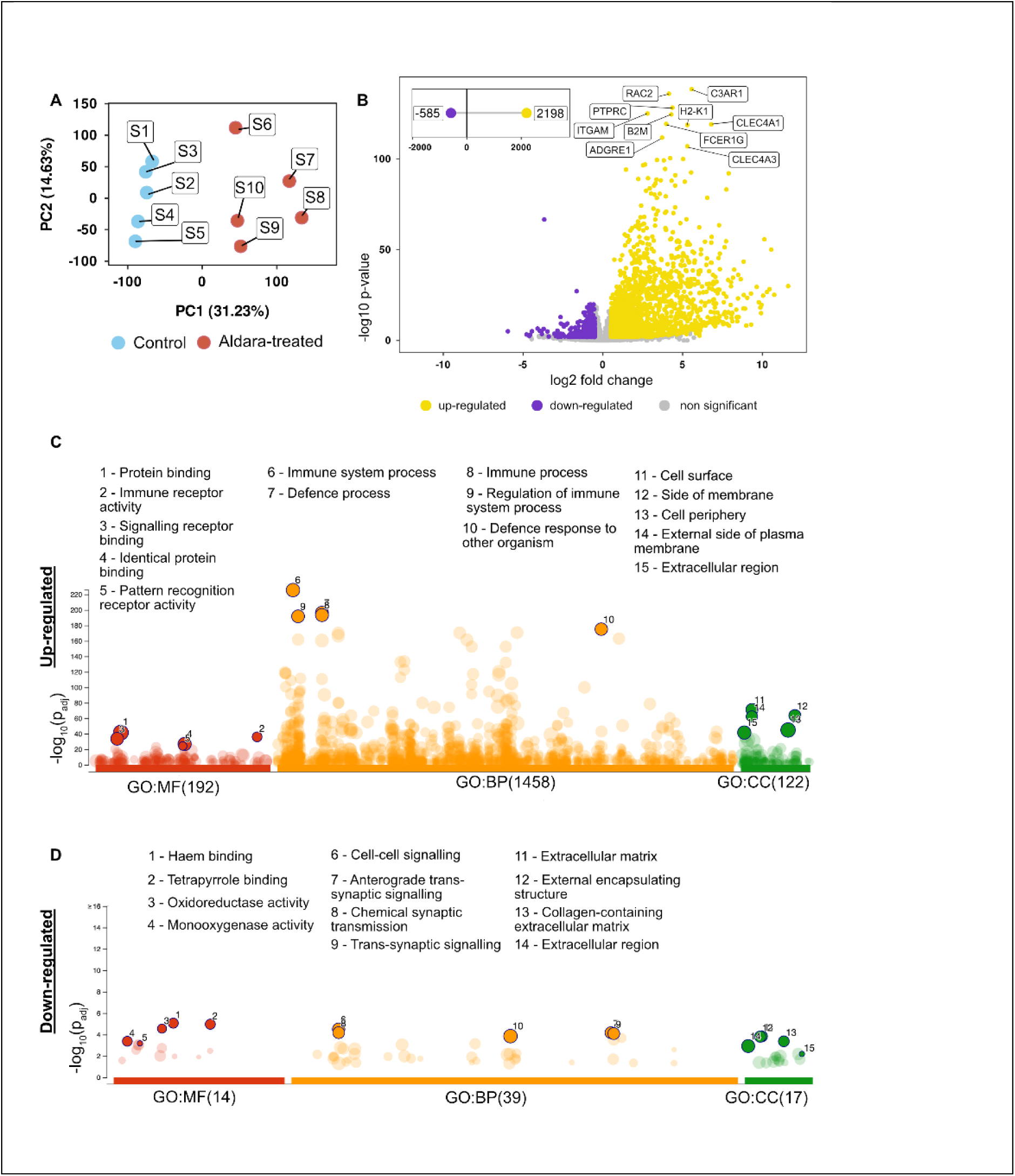
Transcriptome-level changes show a shift in immune-related pathways in the brain following a 3-day inflammatory treatment. **A**, Gene expression data principal component analysis (PCA) scatterplots with each point representing an animal. The percentage of total variation explained by each component is given in the X and Y-axis as appropriate. **B**, Volcano plot for differential expression analysis of Aldara treated samples relative to controls. Significantly differentially expressed genes (p value adjusted <0.05, absolute log_2_ fold > 0.5) are shown in yellow (up-regulated) and purple (down-regulated), with non-significant genes in grey. Inset shows the number of significantly up- and down-regulated genes. **C & D**, Manhattan plots showing enriched pathways in overrepresentation analysis (ORA) for up-regulated (**C**) and down-regulated (**D**) genes. ORA carried out using KEGG, Reactome and WikiPathways databases.

We next carried out an over-representation analysis (ORA) using G:Profiler^29^ to gain a broad understanding of which biological functions were perturbed at the global level in response to Aldara treatment. We ran the ORA using an ordered query to G:Profiler (sorted by log fold change in expression) for all significantly up- and down-regulated genes, run against three Gene Ontology (GO) databases (Molecular Functions [MF], Biological Processes [BP] and

Cellular Components [CC])^30, 31^. Manhattan plots for the ORA for up-regulated and down-regulated gene sets are shown in Figure 1C and 1D, respectively, with full datasets available in supplementary material. Perhaps unsurprisingly given the stimulus, the key upregulated pathways after Aldara treatment centred around immune cell activation, anti-viral and inflammatory responses, cytokine, chemokine and toll-like receptor signalling pathways and those related to microglial activation (Figure 1C). There were substantially fewer downregulated pathways, with the ORA revealing that the main affected pathways were those involved in synaptic signalling and the extracellular matrix.

### Identification of regionally and transcriptionally defined cell clusters following neuroinflammation

The whole brain transcriptome analysis revealed that global changes in transcription in response to 3 days of Aldara treatment matched the immune and neurophysiological changes that we reported previously^6^. We next sought to determine whether there were regionally specific changes in gene expression in brain areas associated with reward and motivation. To this end, we deployed the Nanostring CosM× 1000-plex Mouse Neuroscience RNA panel (for full panel see Supplementary Table 1) to investigate transcriptomic changes in different cell types and brain regions of Aldara- and control-treated mice. We used one anterior hemi-section containing prefrontal cortex and ventral striatum and one posterior hemi-section containing thalamus, amygdala and dorsal hippocampus from each animal (n=4 per group, 16 samples in total). The CosMx Neuroscience Panel is primarily designed to enable exploration of brain-resident cells and neural functions, so contained a relative paucity of immune-related genes. Thus, it was not possible to identify infiltrating immune cells with absolute confidence, nor to parse microglia from infiltrating monocytes. Although the CosMx Mouse Neuroscience Panel contains several microglial markers (e.g. *MerTK, Csf1r, Cx3cr1, Msr1, Hexb* and *Itgax),* many of these are also expressed on infiltrating and border associated myeloid populations^32,33^.

To segment the cell types detectable in our experiment, all samples (both sections from each animal) were integrated for single-cell sequencing-like clustering analysis. This revealed 18 cell clusters, represented on the UMAP in Figure 2A, with the location of cells comprising these clusters shown in representative images in Figure 2D and Supplementary Figure 2. The clusters correspond with the heatmap in Figure 2B that shows unique relative inter-cluster gene expression from the 1000-plex RNA panel. Each row in Figure 2B represents a single gene and is colour-coded to match the UMAP in Figure 2A. A larger version of the heatmap is presented in Supplemental Figure 1 with all gene names added and representative images of 18 clusters and cell counts for posterior slices are on Supplementary Figure 2.

**Figure 2:**
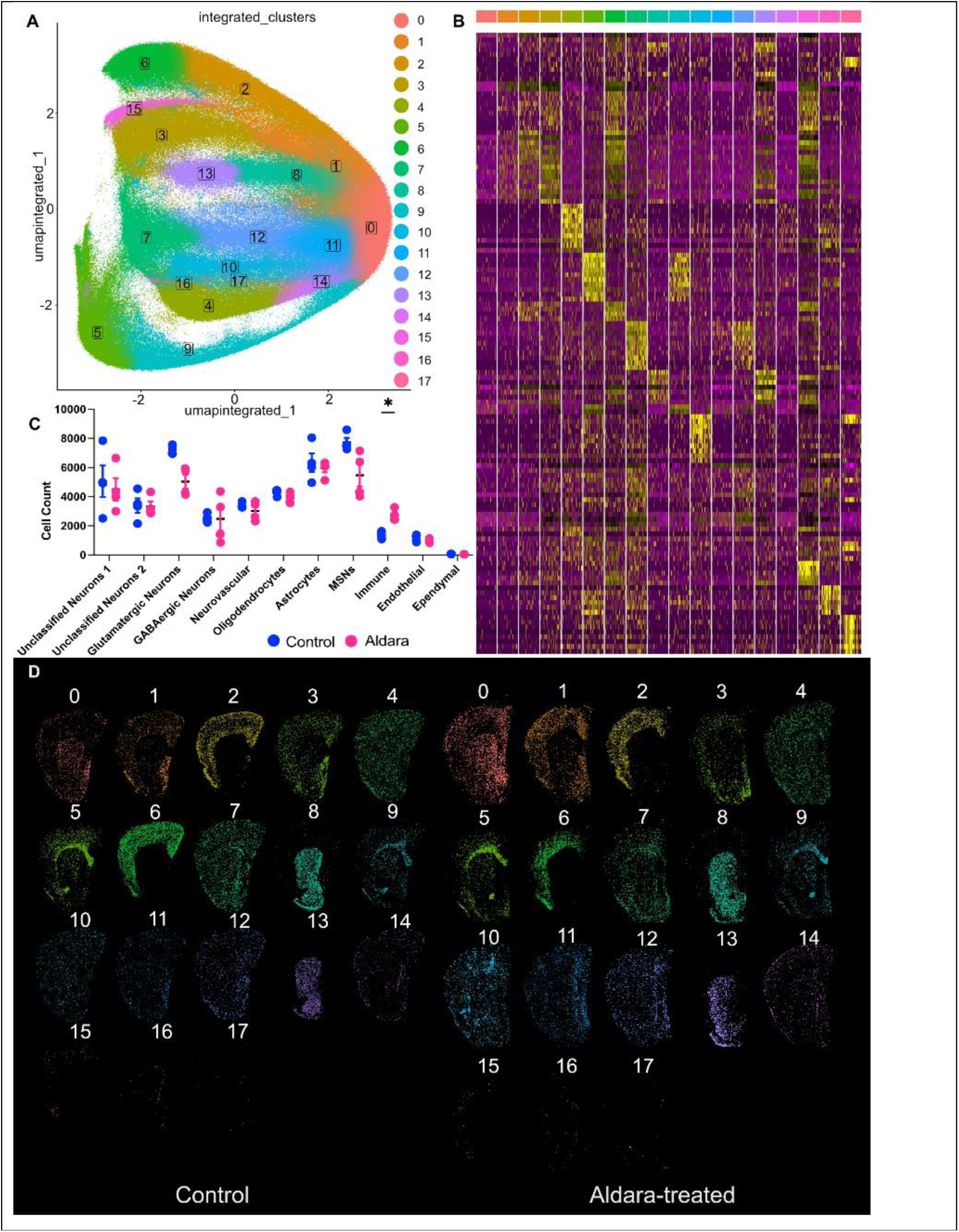
Spatial transcriptomic analysis identified 18 different cell clusters with unique distribution patterns. **A**, An integrated UMAP of all the cells from both control and Aldara-treated samples, showing the distribution of 18 cell clusters. **B**, Heatmap of all clusters showing relative expression of genes. **C**, Cell counts for different cell types found in the anterior slices show a significant increase in the number of immune cells in the Aldara-treated samples. Immune cells control vs Aldara-treated 1367.5 ± 224.9 vs 2720.8 ± 392.7, p = 0.0432 as measured with two-way ANOVA (with Sidak’s multiple comparison test). **D**, Representative images of all clusters and their spatial distributions in control (left) and Aldara-treated (right) anterior brain samples.

Individual cell types were occasionally spread across more than one cluster. Cell type identification was completed by using cluster-specific ‘identifier’ genes coupled with the spatial organisation of the cluster on the coronal tissue sections (Figure 2D). Our analysis parsed 9 unique cell types: glutamatergic neurons, GABAergic neurons, striatal medium spiny neurons (MSNs), astrocytes, oligodendrocytes, endothelial cells, neurovascular unit (NVU; consisting of a mix of endothelial cells and astrocytes), immune cells (inclusive of microglia), ependymal cells and unclassified neurons. The identity of each cluster and the identifier genes are presented in Table 1. While MSNs in the striatum are GABAergic, their transcription profile was sufficiently distinct from other GABAergic neurons to parse these as separate cell types, with their spatial location indeed restricted to striatum (see clusters 8 and 13 of Figure 2D).

To further confirm cell type identification, commonly expressed up-regulated and down-regulated genes across cell types are presented in upset plots (Supplemental Figure 5). For the cases of genes being scientifically unexpected in specific cell types, this may be explained by the limitation of cellular segmentation of the CosMx platform with cellular boundaries potentially displaying overlapping between cell types.

### Astrocytic gene expression changes globally following induction of neuroinflammation

Astrocytes play a dual role as regulators of the neuronal microenvironment and as effectors of immune activation^11^. Our data show that after 3 days of treatment with Aldara, astrocytes increased production of cytokines including TNFα and IL-6 despite displaying no overt changes in morphology^6^. We identified three clusters of astrocytes (clusters 7, 11 and 12) in our samples, providing an opportunity to further explore the effects of Aldara-driven neuroinflammation at the transcriptome level. Astrocytes were identified by the relative inter-cluster upregulation of key astrocytic markers, including *Gpr37l1, Slc1a3, Gja1* and *Aqp4* ^34, 35, 36, 37^. These three clusters were distributed evenly throughout the brain in both anterior (Figure 3A) and posterior (Figure 3B) sections in both control- and Aldara-treated mice.

**Figure 3:**
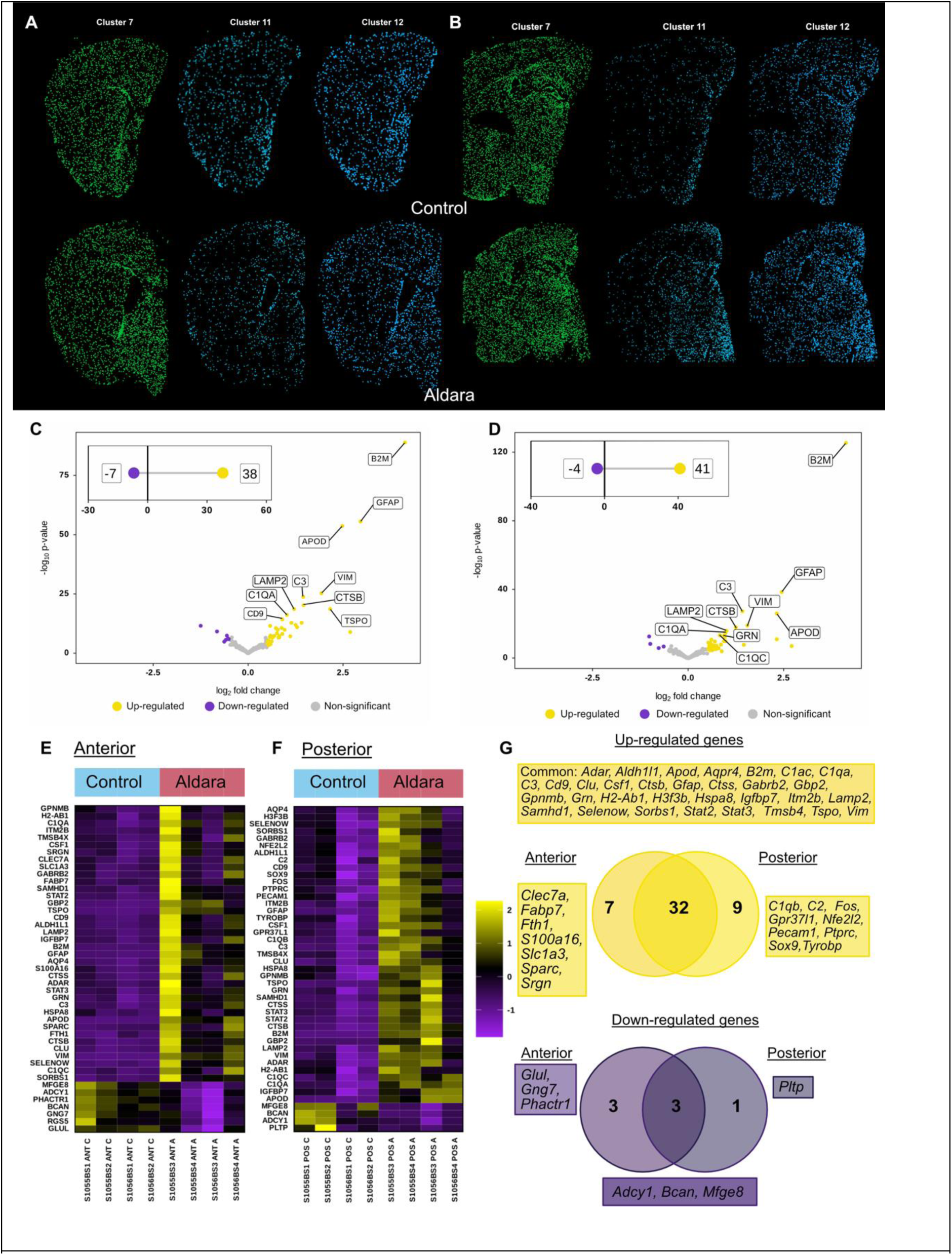
Astrocytic clusters (Cluster 7, 11 and 12) show a large degree of overlap in the up-regulated genes in both posterior and anterior sections. **A**, Representative images of three astrocytic clusters in control (top row) and Aldara-treated conditions (bottom row) in anterior sections, Bregma +1.1-0.8 mm. **B**, Representative images of three astrocytic clusters in control (top row) and Aldara-treated conditions (bottom row) in posterior sections, Bregma -1.0-2.0 mm. **C**, Volcano plot for differential expression analysis of Aldara treated anterior sections relative to controls. Significantly differentially expressed genes are considered by adjusted p value <0.05, absolute log_2_ fold > 0.5, top 10 genes by p value are labelled. Insert shows the number of significantly up- and down-regulated genes. **D**, Volcano plot for differential expression analysis of Aldara treated posterior sections relative to controls. Insert shows the number of significantly up- and down-regulated genes. **E**, Heatmaps of significantly up- and down-regulated genes in Aldara-treated samples as compared with controls. **F,** Overlapping and uniquely upregulated (above) and down-regulated (below) genes of all three astrocytic clusters in anterior and posterior slices.

**Figure 4:**
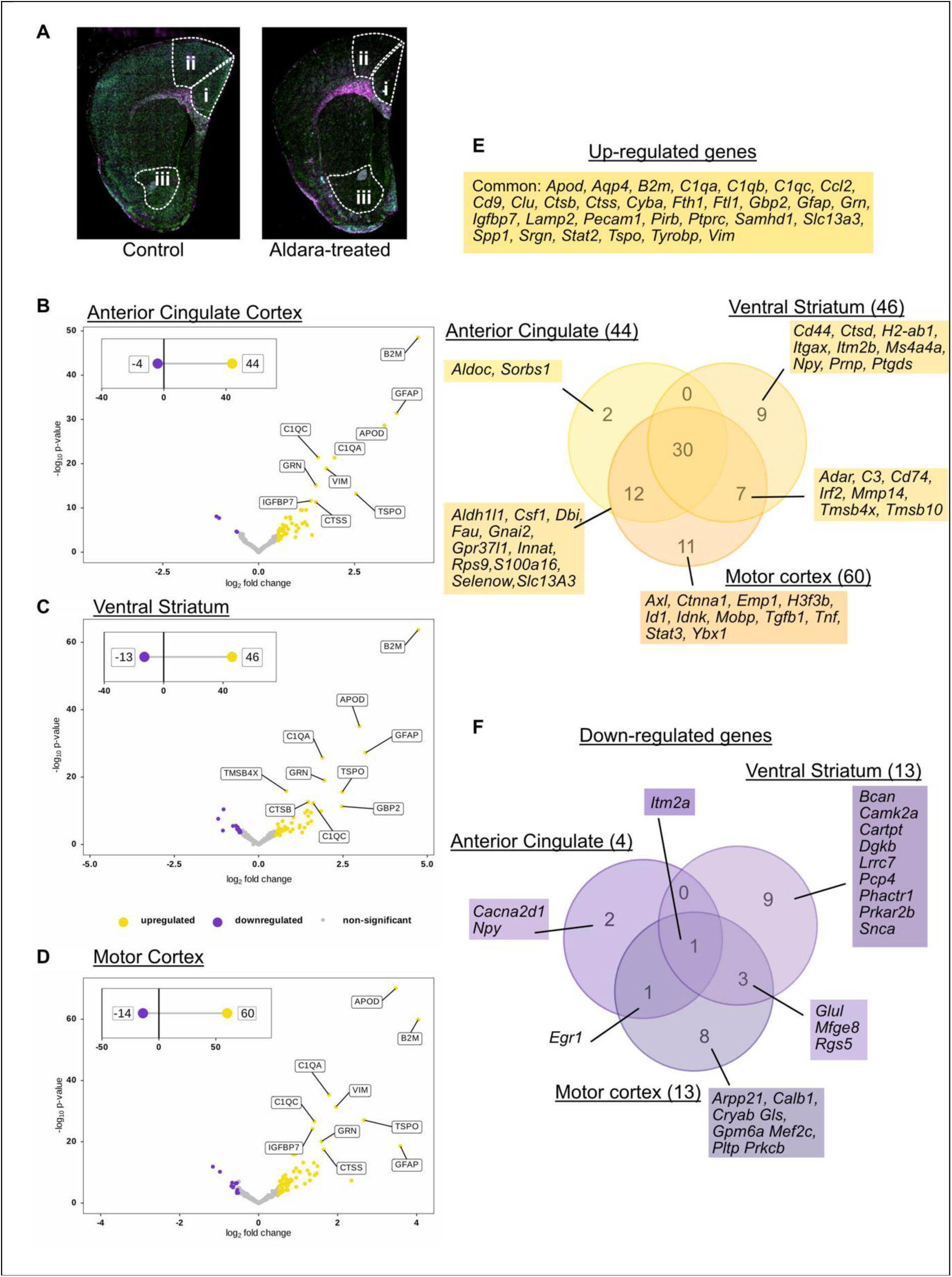
Cluster agnostic approach shows a high degree of overlap in up-regulated genes in anterior cingulate, ventral striatum and motor cortex regions of anterior slices and unique patterns of down-regulated genes. **A**, Representative images of ROI selection in control (left) and Aldara-treated conditions (right) anterior sections, Bregma +1.1 to +0.8 mm. **B-D**, Volcano plot for differential expression analysis of Aldara treated anterior cingulate cortex (**B**), ventral striatum (**C**) and motor cortex (**D**) relative to controls. Insert shows the number of significantly up- and down-regulated genes. Significantly differentially expressed genes are considered by adjusted p value <0.05, absolute log_2_ fold > 0.5, top 10 genes by p value are labelled. **E**, Overlapping and uniquely up-regulated genes in ventral striatum, anterior cingulate and motor cortices. **F**, Overlapping and uniquely down-regulated genes in ventral striatum, anterior cingulate and motor cortices.

The three astrocytic clusters were combined for differential expression analysis using the automated Searchlight2 pipeline^18^. Anterior and posterior tissue samples were analysed separately to ensure that only 1 sample from each animal was used per analysis. PCA analysis revealed two non-overlapping groups for control- and Aldara-treated animals (Supplementary Figure 3A). Volcano plots for both anterior (Figure 3C) and posterior (Figure 3D) sections revealed a similar number of significantly up-regulated and down-regulated genes in the astrocytes from both coronal planes, with 4 to 7 genes downregulated and 38 to 41 genes upregulated. The top 10 significant differentially expressed genes (DEGs) ranked by adjusted p value for anterior sections were *B2m, Gfap, Gbp2, Apod, Tspo, Vim, Ctsb, C3, Stat2* and *Sorbs1.* For posterior sections, the top 10 DEGs were similar: *B2m, Gbp2, Gfap, Tspo, Apod, Vim, Stat2, C3, Ctsb* and *Lamp2.* All significantly DEGs for all anterior and posterior astrocytes are shown in Figures 3E and 3F, respectively. We found that the DEGs between anterior and posterior sections overlapped substantially (Figure 3G). The small number of downregulated genes were involved in neurotransmission and extracellular matrix maintenance.

Commonly upregulated genes included GFAP, the expression of which is known to increase in response to immune and inflammatory stimuli, and genes related to immune function, particularly in the complement family^38, 39^. The changes in gene expression due to Aldara-driven inflammation are discussed below, but a key conclusion from our analysis is that astrocytic response to neuroinflammation occurs globally throughout the brain. Our next aim was to determine whether the impact of neuroinflammation and reactive astrocytes on neuronal function had a similarly global effect, or was regionally specific.

### Neuroinflammation-driven changes in neuronal gene transcription show regional specificity

While astrocytic function is broadly uniform across the whole brain, different regions have substantially different neural architectures in terms of recurrent connectivity, lamination, ratio of glutamatergic to GABAergic neurons, or neuromodulator innervation. We hypothesised that global changes to astrocyte function driven by neuroinflammation could still exert differential effects on local circuitry, with regions associated with reward particularly vulnerable to this disruption. We sought to test this hypothesis by comparing transcriptomic profiles of different brain regions involved in reward with motor cortex, which served as a control region that shared a similar ‘canonical’ architecture to regions such as prefrontal cortex without being involved in reward. Unlike other organs, the brain has a unique 3-dimensional structure with closely interdigitating processes from different cells, such as astrocytic processes enwrapping synapses. Consequently, cell segmentation based on somatic boundaries was unreliable (for example, we observed GFAP expression in almost all identified cell types [supplementary tables 3 and 4]). To compensate for this issue, we adopted a cell-agnostic approach and created regional masks to look at all transcripts within a specific brain region without clustering into specific cell types. This approach allowed us to profit from the advantages of spatial transcriptomics without requiring, e.g., single nucleus sequencing, to confirm specific cell types.

For the anterior sections, our regions of interest were the anterior cingulate cortex (ACC) and ventral striatum (Figure 4) while in posterior sections our regions of interest were the thalamus and amygdala (Figure 5). In response to Aldara treatment, the number of upregulated genes were 44 (ACC; Figure 4B), 46 (ventral striatum; Figure 4C) and 60 (motor cortex; Figure 4D), with 30 of these genes common to all ROIs. The commonly upregulated genes included those involved in immune-mediated processes including complement cascade proteins (*B2m, C1qa, C1qb, Cl1qc, Ctss/b, Grn, Ptprc),* astrocytic reactivity (*Gfap, Vim and Tspo)* and BBB remodelling (*Aqp4, Pecam1* and *Clu). Aldoc* and *Sorbs1* were uniquely upregulated in ACC (Figure 4E)*. Aldoc* encodes for an enzyme in glycolysis in which upregulation of the gene has established links to microglia displaying a shift in glycolytic metabolism in inflammatory tissue environments^40^. *Sorbs1* has been shown to down-regulate proinflammatory cytokines by interfering with the normal function of NF-kB pathway^41^. Ventral striatum was found to have 9 uniquely upregulated genes, relative to controls, in response to the Aldara-driven neuroinflammation (Figure 4E). These genes included those associated with microglia and astrocyte reactivity (*Itgax, Cd44),* BBB remodelling (*Ptdgs),* as well as neuropeptide Y (*Npy).* Uniquely upregulated genes in motor cortex included those involved in pro-inflammatory cytokine signalling (*Tnf)* and cellular proliferation (*Emp1, Id1)*.

**Figure 5:**
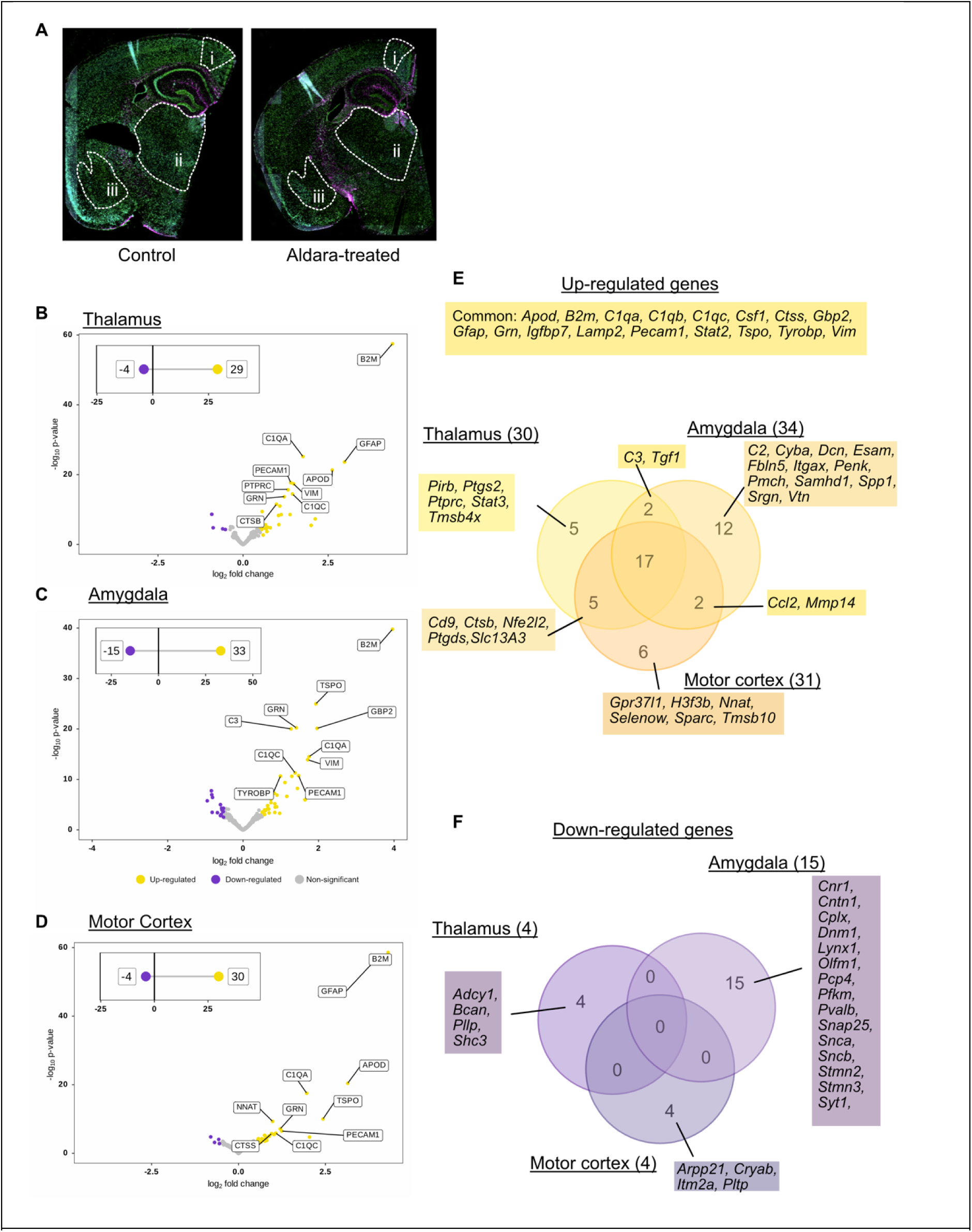
Cluster agnostic approach shows a high degree of overlap in up-regulated genes in thalamus, amygdala and motor cortex regions of posterior slices and unique patterns of down-regulated genes. **A**, Representative images of ROI selection in control (left) and Aldara-treated conditions (right) anterior sections, Bregma range -1.0-2.0 mm. **B-D**, Volcano plot for differential expression analysis of Aldara treated posterior thalamus (**B**), amygdala (**C**) and motor cortex (**D**) relative to controls. Insert shows the number of significantly up- and down-regulated genes. Significantly differentially expressed genes are considered by adjusted p value <0.05, absolute log2 fold > 0.5, top 10 genes by p value are labelled. **E**, Overlapping and uniquely up-regulated genes in thalamus, amygdala and motor cortices. **F,** Overlapping and uniquely down-regulated genes in thalamus, amygdala and motor cortices.

Relatively fewer genes were found to be significantly downregulated in anterior sections, with 4 (ACC; Figure 4B), 13 (ventral striatum; Figure 4C) and 14 (motor cortex; Figure 4D) DEGs detected. Only one gene was commonly downregulated amongst all three ROIs: *Itm2a,* encoding for integral membrane protein 2A that, in the brain, is primarily expressed in the vascular cells^42^. In ACC, two genes were significantly and uniquely downregulated: the voltage-gated calcium channel subunit *Cacna2d1* and the neuropeptide gene *Npy.* In ventral striatum, 9 genes were significantly downregulated, particularly those associated with glutamatergic neurons (*Camk2a*), synapses (*Lrrc7*; *Prkar2b)*; *Snca,* encoding ɑ-synuclein, and other aspects of synaptic transmission (*Dkgb; Pcp4; phactr1*). Motor cortex had 8 uniquely downregulated eight genes which again mainly featured neuronal signalling and transmission genes including *Arpp21, Gls* and *Prkcb.* Additionally, the motor cortex features some neuroprotection-suggestive genes including *Cryab* and *Pltp*.

For posterior sections, our regions of interest were thalamus and amygdala, again using motor cortex as a control region (Figure 5A). In response to Aldara treatment, the number of upregulated genes were 29 (thalamus; Figure 5B), 33 (amygdala; Figure 5C) and 30 (motor cortex; Figure 5D), with 17 of these genes common to all ROIs. The commonly upregulated genes amongst posterior ROIs were similar to those in the anterior sections, plus the addition of *Csf1,* which encodes colony stimulating factor 1, a gene essential for homeostasis and survival of microglia and tissue-resident macrophages^43^. Unique DEGs in the thalamus were related to immune biology including cellular infiltration (*Ptprc)* and anti-inflammatory processes (*Pirb* and *Tmsb4x).* In the amygdala, uniquely upregulated DEGs included those involved in complement cascade (*C2),* endothelial cell functions (*Esam),* extracellular matrix remodelling (*Fbln5* and *Dcn)* and microglia-mediated neuroinflammation (*Itgax* and *Srgn).* The gene encoding the neuropeptide proenkephalin, which is involved in stress responses^39^, was also upregulated.. In motor cortex, uniquely upregulated DEGs encoded proteins involved in anti-oxidative stress mechanisms (*Selenow),* astrocytic neuroprotection (*Gpr37l1)* and anti-inflammatory mechanisms (*Nnat)*.

As with anterior regions, in posterior sections we also observed fewer significantly downregulated genes than those being upregulated (Figure 5E), with 4 (thalamus; Figure 5B), 15 (amygdala; Figure 5C) and 14 (motor cortex; Figure 5D) DEGs detected. There was no overlap in genes significantly downregulated in response to Aldara treatment amongst these three brain regions. In thalamus, significantly downregulated genes were those involved in perineuronal nets (*Bcan*), myelin function (*Pllp*) or neurotransmission (*Adcy1, Shc3*). In the amygdala, downregulated genes included those encoding cannabinoid receptor 1 (*Cnr1*), synapse formation and function (including *Snca, Sncb, Dnm1, Snap25, Lynx1*). Interestingly, the gene encoding parvalbumin (*Pvalb*) was also specifically downregulated in amygdala, suggesting perturbations in interneuron networks. In motor cortex, genes involved in both pro-/anti-inflammatory processes (*Itm2a, Pltp* and *Cryab)* were uniquely downregulated, along with *Arpp21*, encoding a protein involved in regulating dendritic complexity.

## Discussion

We previously found that topical treatment of Aldara drives a combined peripheral and central inflammatory response in mice^13, 14, 15^ that is associated with immune cell infiltration into the brain, cytokine production by resident glial cells, perturbations of thalamostriatal signalling and anhedonia-like behaviour^6^. In the present study, we used a combination of whole-brain bulk RNAseq and spatial transcriptomics using the CosMx platform to study global and local changes in gene expression in response to 3 consecutive days of Aldara treatment, which corresponds to peak neuroinflammation. Overall, changes in gene expression were indicative of a global inflammatory response, confirming our previous results at the transcript level, while also suggesting that this global inflammation results in regionally-specific reductions in gene expression related to neurotransmission and synaptic function.

### Brain-wide transcriptomic changes in response to Aldara treatment

Whole-brain RNA sequencing provided an unbiased measure of the transcriptional response to Aldara. There were marked differences between Aldara brains and those of the controls. Among the most enriched pathways for upregulated gene sets were the innate immune response, type II interferon response, chemokine and TLR signalling pathway which concurs with our previous data on Aldara brain responses^13, 14^. Transcriptomic changes also revealed altered expression of cytokines and chemokines in Aldara-treated mice, relative to controls, that were consistent with a proinflammatory response. For example, TNFα, IL6, IL1β and IFNƴ were all upregulated. These data replicate those that we have previously reported^13, 14, 28^ using methods such as flow cytometry.

We found an upregulation of chemokines signalling pathways which highlight chemokines as being likely drivers of immune cell recruitment during neuroinflammation. Immune system gene expression was confirmed by infiltration of B cells, T cells, Monocytes, NK and NKT cells via flow cytometry in our previous study^6^. Cell signature pathways were also upregulated at transcriptional level, and we found a distinct Th1/Th17 T cell response in this model^6^. Several key microglial activation pathways were among the most upregulated including: microglia phagocytic pathway, DAP12, which along with TREM2 is the principal regulator driving microglia to reactive state and regulates microglial cytokine production^44^ . *Tyrobp* is expressed on microglia and serves as an adaptor for immune receptors including *TREM2* and is robustly upregulated in early transition of microglia to reactive phenotypes^45^.

The most enriched pathways for downregulated genes were those related to neurotransmission, which was consistent with our electrophysiological findings of impaired thalamostriatal signalling reported previously^6^. Overall, these global transcriptional changes indicate a clear neuroinflammatory response focused on IFNγ and TNFa as well as an immune cell response, notably T cell, to the Aldara stimulus. For both downregulated and upregulated gene pathways, data from the present transcriptomic study were highly complementary to those arising from prior neurobiological and immunological experiments. While the whole brain transcriptome approach provides a useful measure of large-scale changes in the brain, the spatial approach was required to provide a more nuanced measure of changes to different neural circuits.

### Astrocytic response to neuroinflammation

As discussed earlier, astrocytes have a key role in regulating both immune and neural functions within the brain, providing an important bridge between these two systems^11^. The astrocytic response appeared to be uniform across the entire brain, suggesting that immune activation in the brain has a similar effect throughout. This is consistent with our previous findings showing reactive microglia in all brain regions studied, and a uniform distribution of T cells into the brain parenchyma^6^. In astrocytes, significantly upregulated genes included those suggesting an astrocytic response to inflammation, namely *Gbp2*, *Gfap, Tspo*. TSPO is an outer mitochondrial membrane protein that is known to be upregulated during neuroinflammation as a sign of mitochondria involvement^46^. Astrocytic contribution to classic immune activation processes is suggested by upregulation of key genes involved in these pathways. Upregulated genes within the identified astrocyte population which have established links to antigen presentation include *B2m, H2-Ab1* and *Ctss.* This increase in antigen presentation-related genes may indicate classical immune activation between resident and infiltrating immune populations, however, *B2m* has also been established as an essential component for astrogliosis *in vitro*^47^.

One of the most striking features of astrocytic gene expression was the upregulation of complement genes, particularly those encoding C1q. The complement cascade is an important effector of synaptic elimination in the CNS in a process mediated by astrocytes^48^. While synapse elimination is typically associated with microglial activation, astrocytes have been shown to phagocytose adult synapses during normal homeostasis^49^ and emerging findings suggest that astrocytes might actually be more likely than microglia to prune synapses during pathological conditions^50^. While we have not been able to demonstrate direct evidence of astrocytic pruning of synapses in our model, these data combined with downregulation in synapse-related genes provide consistent evidence that would support this interpretation.

Within astrocytes, comparatively fewer genes were significantly downregulated in response to Aldara-driven inflammation. Across all sections, three genes were commonly downregulated: those encoding adenylyl cyclase, brevican (a component of perineuronal nets) and MFG-E8, which is regulates phagocytosis of synapses by astrocytes and microglia ^51^. The downregulation of the adenylyl cyclase gene is interesting as we also saw an increase in the expression of the *Gababr2* in astrocytes, which encodes for the B2 subunit of GABA_B_ receptor, which functions by inhibiting adenylyl cyclase via Gi/o^52^. GABA_B_ receptor activation on astrocytes can promote an anti-inflammatory inflammatory phenotype in astrocytes^53^ through inhibition of the NFkB and P38 MAP kinase inflammatory pathways^54^. In anterior sections containing a large amount of striatum, we saw downregulation of *Glu* (encoding glutamine synthetase), *Gng7* (also related to adenylyl cyclase function^55^ and Phactr1, related to protein phosphorylation^56^.

Our data also suggest additional neuroprotective mechanisms that may be activated in astrocytes. Clusterin, coded for by *Clu*, has been shown to enhance excitatory neurotransmission in preclinical models of Alzheimer’s disease^57 58^, the pathophysiology of which involves inflammation. The glycoprotein encoded by *Gpnmb* has previously been found to hold a neuroprotective role in an astrocyte-specific attenuation of neuroinflammatory environments^59^. Altogether, changes in astrocyte gene expression indicate an increase in expression of genes related to neuroinflammation and a transition to a reactive phenotype, as well as the emergence of anti-inflammatory compensatory mechanisms.

### Neuroinflammatory effects on neural circuitry

We next sought to determine the effects of Aldara-driven neuroinflammation on neural circuit function in specific brain areas. Due to the challenges in accurately delineating the boundaries of neurons and astrocytes with certainty (see later discussion), we examined all transcripts within a particular region and analysed the expression in a manner that was agnostic of cell types. These samples would include a large number of astrocytes, given their ubiquity throughout neural tissue, so it is unsurprising that there was significant upregulation in inflammation-related genes and their pathways, as discussed in the above section. But noting the genes that were uniquely downregulated in specific regions, or that tend to be expressed on neurons, still provides some insight into inflammation-driven changes in circuit function.

We saw a significant increase in neuropeptide (NPY) gene expression in ventral striatum but a decrease in expression in anterior cingulate cortex, two different parts of the reward circuitry. Interestingly, NPY is expressed by some GABAergic neurons in the cortex while it is expressed by non-GABAergic cholinergic interneurons in the striatum^60^, so the effect of this neuropeptide likely has region-specific effects. In striatum, NPY expression can increase in response to dopamine depletion^61^ while in ACC, a decrease in NPY expression can result from chronic stress^62^. As described in the results, in ventral striatum, we saw downregulation of several genes involved in synaptic transmission or homeostasis. We saw a significant downregulation of *Camk2a*, a ubiquitous marker of glutamatergic neurons. While there are no glutamatergic neuronal cell bodies in ventral striatum, there are an abundance of excitatory axon terminals arriving from both cortex and thalamus^63^. In Drosophila, it was recently shown that *Camk2a* mRNA is locally translated in axons in a mechanism required for memory formation^64^, although it is not yet clear whether such a mechanism operates in the mammalian brain. However, the existence of local mRNA transport to, and translation in, axons has become widely accepted^65^. In both amygdala and ventral striatum, we saw downregulation of genes associated with synaptic release, such as *Snca* and *Pcp4* amongst many others, suggesting reductions in inputs to these regions. This provides mechanistic insight to some of the mechanisms that may underlie the significant impairment of the ability of thalamic axons to excite ventral striatum MSNs that we reported previously^6^.

One of the interesting findings of our previous work on behavioural consequences of Aldara-driven neuroinflammation was the presence of an anhedonia-like phenotype without any obvious anxiety behaviours^6^. In amygdala, there was unique upregulation of the gene encoding proenkephalin; others have shown that overexpression of proenkephalin in amygdala enhances the anxiolytic effect of benzodiazepines^66^, and knockdown of this protein in the amygdala is sufficient to induce anxiety-like behaviours in rats otherwise resistant to the induction of chronic unpredictable stress^67^. The upregulation of the gene encoding proenkephalin may underlie the lack of anxiety-like behaviours in Aldara-treated mice. However, we also saw a reduction in the expression *Cnr1*, encoding the cannabinoid receptor 1, in amygdala. Others have shown that reducing expression of this gene in amygdala can increase anxiety or depression-like behaviours in mice^68^ and marmosets^69^. These contradictory mechanisms highlight the challenges in over-attributing specific functions to individual genes in transcriptomics datasets, and the need for multiple corroborating pieces of evidence from the same biological pathways.

### Limitations of spatial transcriptomics approach in neural tissue

The main limitation in applying the CosMx spatial transcriptomics method to brain tissue is that the brain has an incredibly complex morphology with processes from different neurons and astrocytes located within tens of nanometres of each other^70^. A single cubic micrometre could contain processes from dozens of cells. Given that local mRNA translation occurs in axons^65^, dendrites^71^ and astrocytic processes^72^, and that we were imaging transcripts in 2 dimensions on tissue sections that were 6 µm thick, it is simply not possible to definitively assign any individual transcript to a specific cell with a high degree of certainty. This may explain the finding of upregulation of the primarily astrocyte-specific gene *Gfap* amongst all identified cell types, when this gene is typically associated with astrocytes. Similarly, we saw high expression of C1q that was attributed to astrocytes in our analysis, although C1q is typically expressed by neurons when being used to tag synapses for elimination^49^. The approach used in our study alone would be unable to confirm whether the C1q mRNA detected in our study was produced by astrocytes or located in nearby neuronal processes and a technique such as combined electron microscopy *in situ* hybridisation^73^ would be required to confirm.

An additional limitation with the CosMx is the size (1000 genes) of the mouse neuroscience panel. While allowing good characterisation of neuronal function, it was more limited for studying brain-immune interactions. An expansion of the panel or coupling the spatial transcriptomics approach with a whole transcriptomic single nuclear sequencing platform would strengthen confidence in cellular characterisation and discovery biology.

### Concluding remarks

In this study, we have employed a spatial transcriptomics method to further understand the changes occurring in the brain in response to Aldara-driven neuroinflammation. Despite challenges created by the complex structure of the brain, our study has shown that a global inflammatory stimulus throughout the brain can evoke substantially different changes in different neural circuits. The next challenge is to identify the links between neurobiology and immune function that are unique to different brain regions, so that we can understand why some circuits seem particularly vulnerable to inflammatory environments and perhaps identify novel targets to treat inflammation-associated depression.

## Acknowledgements

We gratefully acknowledge the support of the Inger M Lilya and George Simpson Biological Psychiatry Scholarships. The authors gratefully acknowledge the Glasgow Imaging Facility for their support and assurance in this work. The authors gratefully acknowledge the Flow Core Facility of the University of Glasgow for their support and assistance in this work. For the purpose of open access, the authors have applied a Creative Commons Attribution (CC-BY) licence to any Author Accepted Manuscript version arising from this submission.

