## Supplementary material for "Neuroinflammation driven by TLR7 activation in mice results in a global inflammatory response driving circuit-specific changes in neuronal gene expression"

**Supplementary table 1. CosMx 1000-plex panel list of genes.**

| Gene | Gene | Gene | Gene | Gene | Gene | Gene | Gene |
|---|---|---|---|---|---|---|---|
| 4930486L24Rik | Ambp | Avp | Calb2 | Chrna7 | Ctnna2 | Dock3 | Fbxw7 |
| 6330403K07Rik | Ambra1 | Avpr1a | Calca | Ckb | Ctnnb1 | Dok6 | Fgf1 |
| Abca2 | Anapc16 | Avpr1b | Calcr | Clasp2 | Ctnnd2 | Drd1 | Fgf2 |
| Abi1 | Ano6 | Avpr2 | Calm1 | Cldn5 | Ctsb | Drd2 | Fgfr2 |
| Abi2 | Apba1 | Axl | Calm2 | Clec7a | Ctsd | Drd4 | Flt1 |
| Abl1 | Apc | B2m | Calm3 | Clip1 | Ctss | Dscam | Fnip2 |
| Abl2 | Apela | B3gnt5 | Camk1d | Clstn1 | Ctxn1 | Dscaml1 | Foxo1 |
| Acaca | Aph1b/c | Bace1 | Camk2a | Clu | Cul1 | Dtl | Foxo3 |
| Acacb | Aplnr | Bace2 | Camk2b | Cmip | Cul5 | Dync1h1 | Fpr2 |
| Ace2 | Aplp2 | Bcan | Camk2d | Cnksr2 | Cx3cr1 | Dync1i1 | Fpr3 |
| Acer3 | Apoe | Bcas1 | Camk2g | Cnr1 | Cxcl14 | Dync1i2 | Frs2 |
| Acta2 | App | Bche | Camk2n1 | Cntn1 | Cyfip1 | Dync1li2 | Fth1 |
| Ada | Aqp4 | Bcl2 | Camk4 | Cntn2 | Cyfip2 | Dynll1 | Ftl1 |
| Adam10 | Arhgef7 | Bcl2l1 | Car8 | Cntn4 | Daam1 | Dynll2 | Fus |
| Adam22 | Arih1 | Bdnf | Cartpt | Cntn5 | Daam2 | Eda | Fxyd6 |
| Adar | Arpp21 | Bex1/2 | Casp3 | Col4a5 | Dab1 | Edn1 | Fyn |
| Adcy1 | Arrb1 | Bin1 | Cast | Comt | Dapk1 | Edn2 | Fzd3 |
| Adcy2 | Artn | Birc6 | Cblb | Cop1 | Dapk2 | Edn3 | Gab1 |
| Adcy5 | Aspa | Bmpr1a | Cbln2 | Cox4i1 | Dbi | Ednra | Gab2 |
| Adcy8 | Astn2 | Bmpr1b | Cck | Cox6c | Dcc | Ednrb | Gabbr2 |
| Adcy9 | Atf6 | Bmpr2 | Cckar | Cox7c | Ddt | Eef2k | Gabra2 |
| Add1 | Atg10 | Braf | Cckbr | Cox8a | Deptor | Efna5 | Gabrb1 |
| Adgrg1 | Atg13 | Brwd1 | Ccnf | Cpb2 | Dera | Egfr | Gabrb2 |
| Adgrl2 | Atg2b | Bsg | Cd109 | Cpe | Des | Egr1 | Gabrb3 |
| Adgrl3 | Atg4c | Btrc | Cd14 | Cplx1 | Dgkb | Eif4a2 | Gabrg3 |
| Adgrv1 | Atg5 | C1qb | Cd44 | Cr1l | Dgkg | Eif4g3 | Gad1 |
| Adora1 | Atg7 | C1qc | Cd46 | Creb5 | Dgki | Elmo1 | Gad2 |
| Adora2a | Atm | C1s1/2 | Cd55 | Crebbp | Dicer1 | Emcn | Gal |
| Adra1d | Atp1a2 | C2 | Cd74 | Crh | Dlg1 | Emp1 | Galr1 |
| Adrb2 | Atp1b1 | C3 | Cd81 | Crhr1 | Dlg4 | Entpd2 | Galr2 |

| | | | | | | | |
|---|---|---|---|---|---|---|---|
| Ago2 | Atp2a2 | C5ar1 | Cd9 | Crhr2 | Dlgap2 | Ep300 | Galr3 |
| Ago3 | Atp2b1 | Cab39 | Cdc14b | Crip1 | Dlk1 | Epb41l2 | Gap43 |
| Agt | Atp2b2 | Cab39l | Cdc27 | Crp | Dmd | Epb41l3 | Gas2 |
| Agtr1a | Atp2b4 | Cacna1a | Cdc6 | Cryab | Dnah14 | Epha6 | Gba |
| Agtr1b | Atp6v0a1 | Cacna1b | Cdk6 | Csf1 | Dnah6 | Epn2 | Gbp2 |
| Agtr2 | Atp6v1a | Cacna1c | Cdk8 | Csf1r | Dnah7a/b/c | Esam | Gbp2b |
| Ahcyl1 | Atp6v1h | Cacna1d | Cfh | Csmd3 | Dnajc1 | F13a1 | Gcg |
| Akt1 | Atp8a1 | Cacna1e | Chchd10 | Csnk1a1 | Dner | Fa2h | Gcgr |
| Akt3 | Atpif1 | Cacna2d1 | Chl1 | Csnk2a1 | Dnm1 | Fabp7 | Gdnf |
| Alcam | Atr | Cacna2d3 | Chrm1 | Cspg5 | Dnm1l | Fau | Gfap |
| Aldh1l1 | Atrn | Cacnb2 | Chrm2 | Cst3 | Dnm2 | Fbln5 | Gfod1 |
| Aldoa | Atxn1 | Cacnb4 | Chrm3 | Cst7 | Dnm3 | Fbn1 | Ghr |
| Aldoc | Atxn2 | Cacng3 | Chrm5 | Ctbp2 | Dock1 | Fbxo33 | Gja1 |
| | Auts2 | Calb1 | Chrna4 | Ctnna1 | Dock10 | Fbxw11 | Gjb1 |

| | | | | | | | |
|---|---|---|---|---|---|---|---|
| Glp1r | Gsn | Il1b | Lepr | Mapk14 | Ndrg4 | P2ry12 | Pik3r1 |
| Glp2r | Gstm1 | Il1rap | Lgi2 | Mapk8 | Ndufa10 | P2ry14 | Pink1 |
| Gls | Gucy1a2 | Il1rapl2 | Lgr4 | Mapkap1 | Ndufa13 | P2ry2 | Pip5k1b |
| Glul | H2-Aa | Il6 | Lgr5 | Mapt | Ndufa4 | Pak1 | Pirb |
| Gm2a | H2-Ab1 | Inpp5a | Lifr | Mark3 | Ndufs1 | Pak2 | Pkn2 |
| Gnai1 | H2-Ea | Insr | Lilra5 | Masp1 | Nedd4l | Pak3 | Pla2g4c |
| Gnai2 | H3f3b | Invs | Lilrb4a/b | Mbp | Negr1 | Pard3 | Plcb1 |
| Gnal | H6pd | Ip6k2 | Lpar1 | Mc1r | Nell2 | Park7 | Plcb4 |
| Gnao1 | Hc | Irf2 | Lpar2 | Mc2r | Neo1 | Parn | Plcl1 |
| Gnaq | Hdac4 | Irf6 | Lpar3 | Mc3r | Neu4 | Pbx1 | Pllp |
| Gnas | Hdac9 | Irs2 | Lpar6 | Mc4r | Nf1 | Pcdh15 | Plp1 |
| Gnb1 | Hecw1 | Itch | Lpin1 | Mc5r | Nfasc | Pcp2 | Plpp3 |
| Gng7 | Herc1 | Itga2 | Lpl | Mcm5 | Nfe2l2 | Pcp4 | Plppr1 |
| Gphn | Herc4 | Itga8 | Lrp2 | Mcu | Nfkb1 | Pcsk1n | Pltp |
| Gpi1 | Hexb | Itga9 | Lrp6 | Mdga2 | Nlgn4l | Pcsk5 | Plxna4 |
| Gpm6a | Higd1b | Itgav | Lrrc4c | Mdh1 | Nlk | Pcsk6 | Pmch |
| Gpm6b | Hip1 | Itgax | Lrrc7 | Mef2a | Nnat | Pde10a | Polr2f |
| Gpnmb | Hipk2 | Itgb8 | Lrrk2 | Mef2c | Nos1 | Pde1a | Pomc |
| Gpr158 | Hmgb1 | Itm2a | Lrrtm3 | Meg3 | Npy | Pde1c | Pou2f1 |
| Gpr17 | Hnrnpa2b1 | Itm2b | Lsr | Megf11 | Npy1r | Pde3b | Ppargc1a |
| Gpr37 | Homer1 | Itm2c | Ltbp1 | Mertk | Npy2r | Pde4d | Ppfia2 |
| Gpr37l1 | Hsp90aa1 | Itpr1 | Lynx1 | Mfge8 | Nr3c1 | Pde7b | Ppp1r13b |
| Grb10 | Hsp90ab1 | Itpr2 | Lyz1/2 | Mib1 | Nrcam | Pde8a | Ppp1r1b |
| Grb2 | Hspa1a | Jak1 | Mag | Mid1 | Nrf1 | Pdgfd | Ppp2r2a |

| | | | | | | | |
|---|---|---|---|---|---|---|---|
| Gria1 | Hspa1b | Jak2 | Maged1 | Mllt11 | Nrg1 | Pdgfra | Ppp2r2c |
| Gria2 | Hspa4l | Jam3 | Magi1 | Mobp | Nrg2 | Pdyn | Ppp2r3a |
| Gria3 | Hspa8 | Jun | Malat1 | Mog | Nrgn | Pecam 1 | Ppp2r5c |
| Gria4 | Hsph1 | Kalrn | Maml2 | Msr1 | Nrsn1 | Peli2 | Ppp2r5e |
| Grid1 | Htr1a | Kat2b | Maml3 | Mt1 | Nsg1 | Penk | Ppp3ca |
| Grid2 | Htr2a | Kcna2 | Man1a | Mt3 | Ntm | Pfkl | Ppp3cb |
| Grik1 | Htr2c | Kcnd2 | Man1a2 | Mtmr3 | Ntrk2 | Pfkm | Ppp3cc |
| Grik2 | Htr3a | Kcnj3 | Man1c1 | Myl6 | Ntrk3 | Pfkp | Prex1 |
| Grin1 | Htr6 | Kcnq3 | Man2a1 | Myl9 | Ntsr2 | Pgd | Prickle1 |
| Grin2a | Htt | Kcnq5 | Maoa | Myo6 | Nup160 | Pgls | Prickle2 |
| Grin2b | Iapp | Kdm4a | Maob | Myrf | Nup214 | Pgm1 | Prkacb |
| Grip1 | Icmt | Kidins22 0 | Map1b | Nap1l5 | Olfm1 | Phactr1 | Prkag2 |
| Grk3 | Idnk | Kif5c | Map2k1 | Nav3 | Olig1 | Phc2 | Prkar1b |
| Grm1 | Ifnar1 | Kirrel3 | Map2k4 | Nbea | Olig2 | Phc3 | Prkar2a |
| Grm3 | Ifngr1 | Ksr2 | Map2k5 | Ncam1 | Opa1 | Phkb | Prkar2b |
| Grm5 | Ifngr2 | Lama2 | Map2k6 | Nckap1 | Oprd1 | Phlpp1 | Prkca |
| Grm7 | Ift88 | Lamc2 | Map3k5 | Ncor1 | Oprk1 | Pias1 | Prkcb |
| Grm8 | Igf1r | Lamp2 | Map4k4 | Ncor2 | Oprm1 | Pik3c3 | Prkce |
| Grn | Igfbp7 | Lancl1 | Mapk1 | Ndrg1 | P2rx4 | Pik3ca | Prkcq |
| Gsk3b | Il10 | Lep | Mapk10 | Ndrg2 | P2rx7 | Pik3cb | Prkg1 |
| Prkn | Rbpj | S100a16 | Slc2a13 | Sspo | Thy1 | Ube2k | Znrf3 |
| Prnp | Reln | S100b | Slc32a1 | Sst | Tiam1 | Ube2r2 | Zwint |
| Pros1 | Rgl1 | Sall1 | Slc38a9 | Sstr1 | Timp2 | Ube3c | |
| Prps2 | Rgn | Samhd1 | Slc39a1 0 | Sstr2 | Tjp1 | Ube4b | |
| Psap | Rgs12 | Sbf2 | Slc39a1 1 | Sstr3 | Tkt | Ubr5 | |
| Psen1 | Rgs5 | Scd2 | Slc44a1 | Sstr4 | Tle4 | Uggt2 | |
| Psma1 | Rgs6 | Scg2 | Slc4a4 | St3gal6 | Tlr4 | Ugt8a | |
| Psmd1 | Rgs7 | Scg5 | Slc4a8 | Stat2 | Tmem119 | Ulk2 | |
| Psmd1 4 | Rhobtb1 | Scn2a | Slc6a1 | Stat3 | Tmsb10 | Unc5c | |
| Psme4 | Rictor | Scn8a | Slc6a3 | Stk3 | Tmsb4x | Unc5d | |
| Pten | Rims1 | Sec24b | Slc8a1 | Stmn2 | Tnc | Uqcr10 | |
| Ptgds | Rims2 | Selenow | Slco3a1 | Stmn3 | Tnf | Uqcrq | |
| Ptgs2 | Rit2 | Selplg | Slit2 | Sulf2 | Tnik | Ush1g | |
| Ptk2 | Rnf144a | Sem1 | Sln | Sv2a | Tnks | Usp15 | |
| Ptn | Robo1 | Sema5a | Smad2 | Syn2 | Tnr | Usp34 | |
| Ptpn1 | Robo2 | Sema6d | Smad3 | Syn3 | Tnrc6a | Usp9x | |
| Ptpn11 | Rock1 | Serinc3 | Smurf1 | Synpr | Tnrc6b | Uvrag | |
| Ptpn13 | Rock2 | Serinc5 | Snap25 | Syp | Tnrc6c | Vav3 | |
| Ptpn4 | Ror1 | Serpine2 | Snca | Syt1 | Tph2 | Vcan | |
| Ptprg | Rora | Serping1 | Sncb | Syt11 | Traf3 | Vegfa | |
| Ptprj | Rpe | Setx | Snhg11 | Tab2 | Trem2 | Vim | |
| Ptpro | Rpia | Sgk1 | Snn | Tac1 | Trim2 | Vip | |

| | | | | | | |
|---|---|---|---|---|---|---|
| Ptprr | Rpl10 | Sgk3 | Snrpn | Tacr1 | Trim33 | Vipr1 |
| Ptprs | Rpl13a | Sh3gl2 | Sod1 | Tacr2 | Trim62 | Vipr2 |
| Ptprz1 | Rpl14 | Sh3gl3 | Sorbs1 | Tacr3 | Trio | Vmp1 |
| Pvalb | Rpl32 | Sh3tc2 | Sorl1 | Tagln | Trip12 | Vtn |
| Rab1a | Rpl37 | Shc3 | Sort1 | Taldo1 | Trp53 | Wac |
| Rab31 | Rplp0 | Shprh | Sos1 | Tank | Trpm7 | Wasf1 |
| Rab6a | Rps14 | Shtn1 | Sos2 | Taok3 | Trpv1 | Wdr41 |
| Rab7 | Rps27 | Sil1 | Sox5 | Tardbp | Tspan3 | Wdr59 |
| Raf1 | Rps4x | Sipa1l1 | Sox6 | Tbl1x | Tspan5 | Wscd1 |
| Ramp1 | Rps5 | Skil | Sox9 | Tbl1xr1 | Tspan7 | Wwp1 |
| Ranbp2 | Rps6ka2 | Slc10a6 | Spag9 | Tcf12 | Tspo | Wwp2 |
| Rap1a | Rps6ka3 | Slc11a2 | Sparc | Tcf4 | Ttc3 | Xist |
| Rapgef 1 | Rps6ka5 | Slc12a2 | Sparcl1 | Tcf7l2 | Ttr | Xpo1 |
| Rapgef 2 | Rps9 | Slc13a3 | Spg11 | Tenm2 | Tuba1a/b/ c | Ybx1 |
| Rasa1 | Rptor | Slc17a7 | Spock1 | Tenm4 | Tubb5 | Ywhae |
| Rasal2 | Rspo2 | Slc18a2 | Spp1 | Terf2ip | Tusc3 | Ywhag |
| Rasgrf2 | Rtn1 | Slc1a2 | Spp2 | Tfdp2 | Tyrobp | Ywhaq |
| Rb1 | Rtn3 | Slc1a3 | Sptan1 | Tgfb1 | Ube2e1 | Ywhaz |
| Rb1cc1 | Rtn4 | Slc24a3 | Sptbn1 | Tgfb2 | Ube2e2 | Zbtb20 |
| Rbfox3 | Runx1 | Slc25a12 | Sptbn4 | Tgfbr1 | Ube2e3 | Zeb1 |
| Rbks | Ryr2 | Slc25a4 | Srgn | Thrb | Ube2g1 | Zfyve1 6 |
| Rbp4 | Ryr3 | Slc2a1 | Ssh2 | Thsd4 | Ube2h | Zfyve9 |

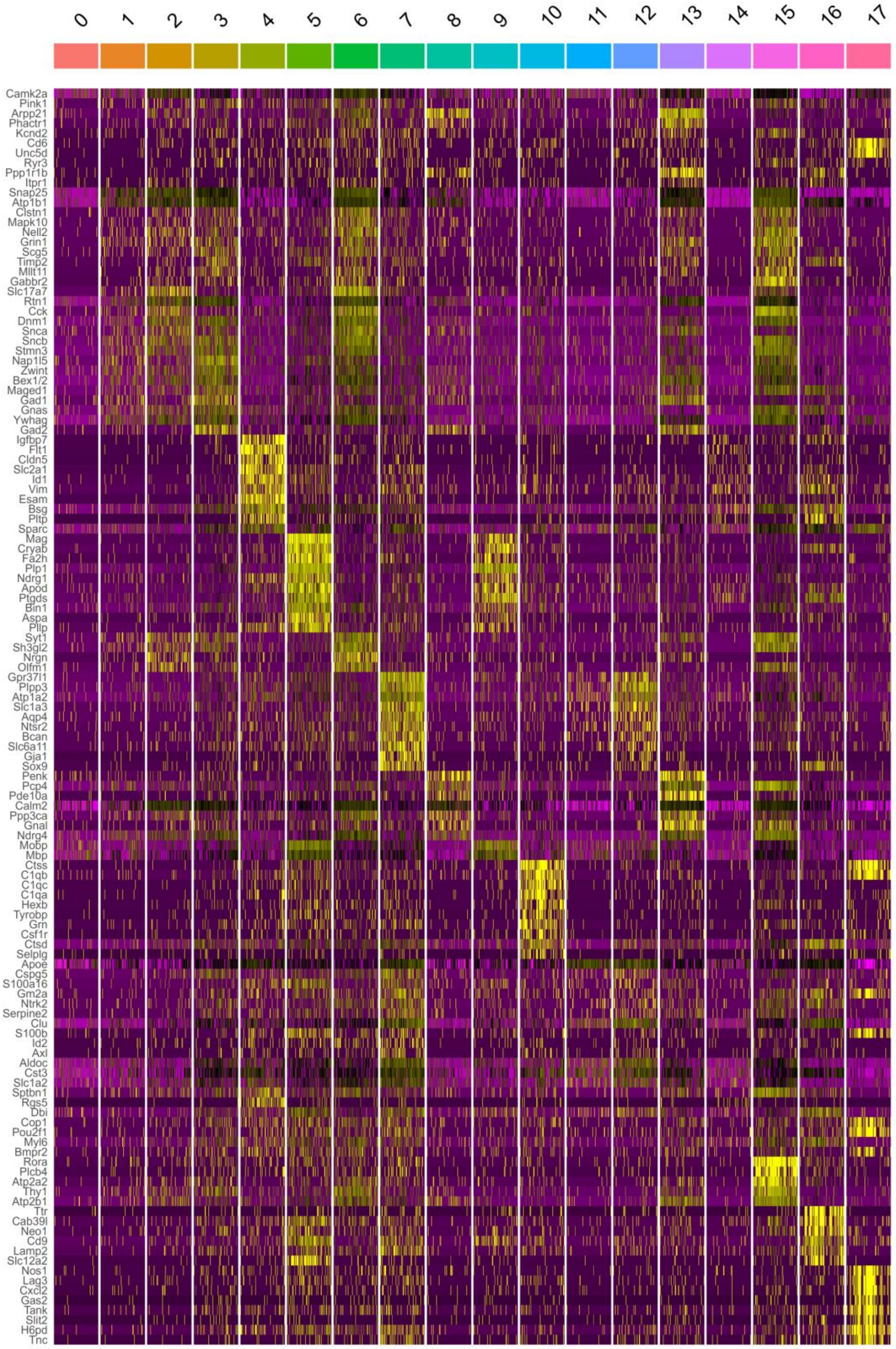

**Supplementary Figure 1: Inter-cluster heatmap of gene expression for 18 identified CosMx clusters with relevant cluster marker genes.** Yellow indicates relative increased gene expression and purple indicates relative decreased gene expression.

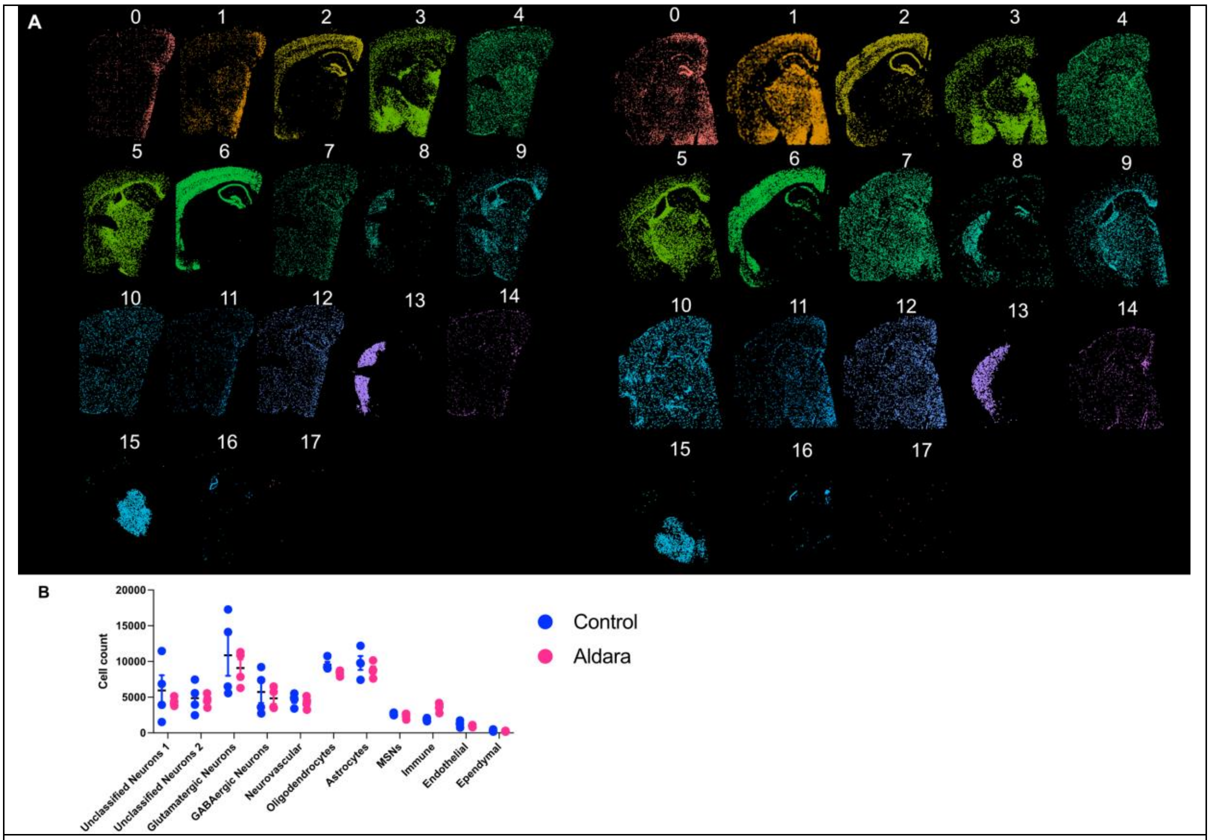

**Supplementary Figure 2: A,** Representative clusters in control (left) and Aldara-treated (right) conditions in posterior sections. **B**, Cell counts for different cell types in posterior sections, no significant differences were found with a two-way ANOVA (with Sidak's multiple comparison test).

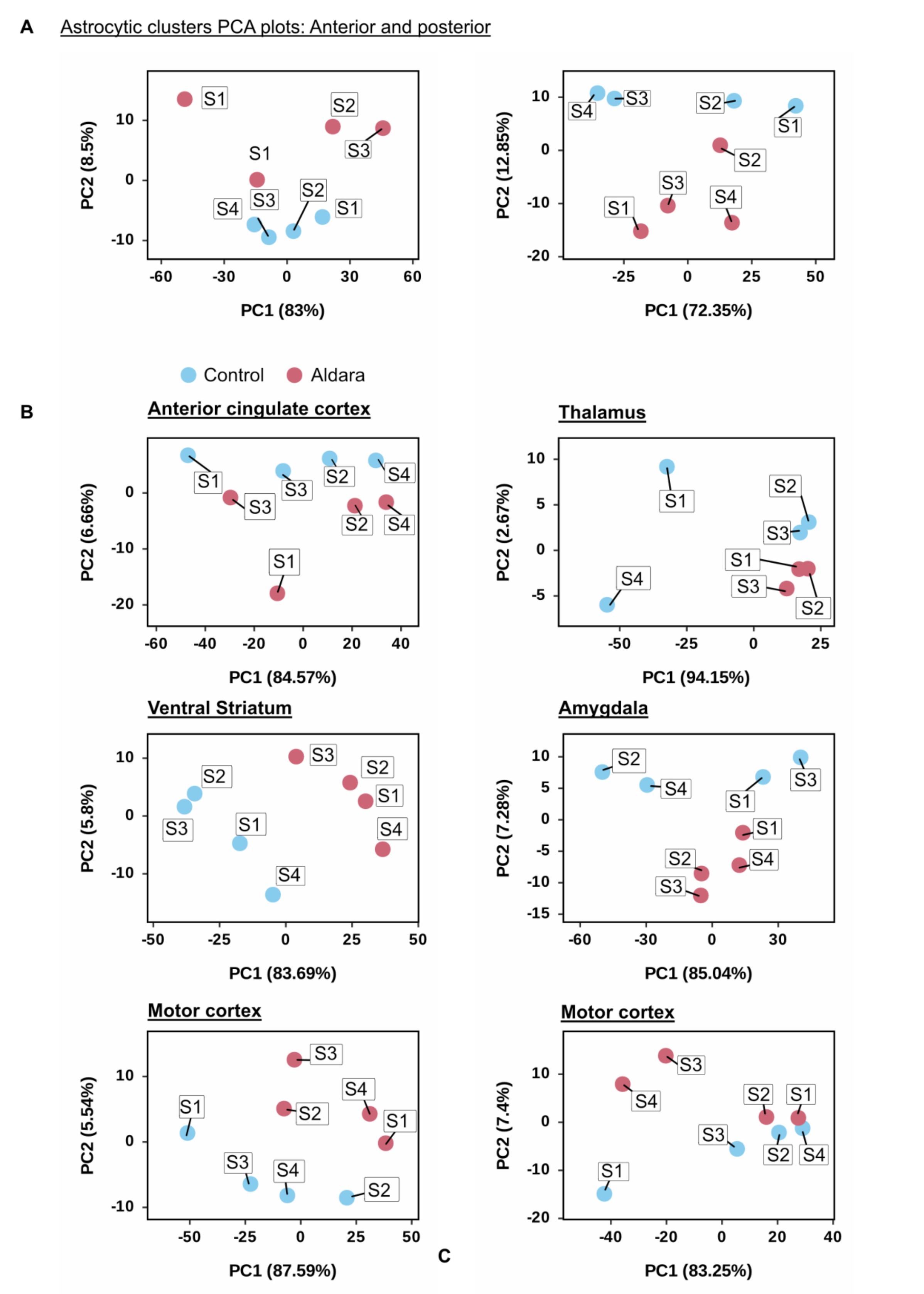

**Supplementary Figure 3: A,** PCA plots for CosMx: astrocytic clusters (clusters 7, 11 and 12) anterior (left) and posterior (right). **B**, PCA plots for regions of interest from anterior sections (left column) and posterior section (right column).

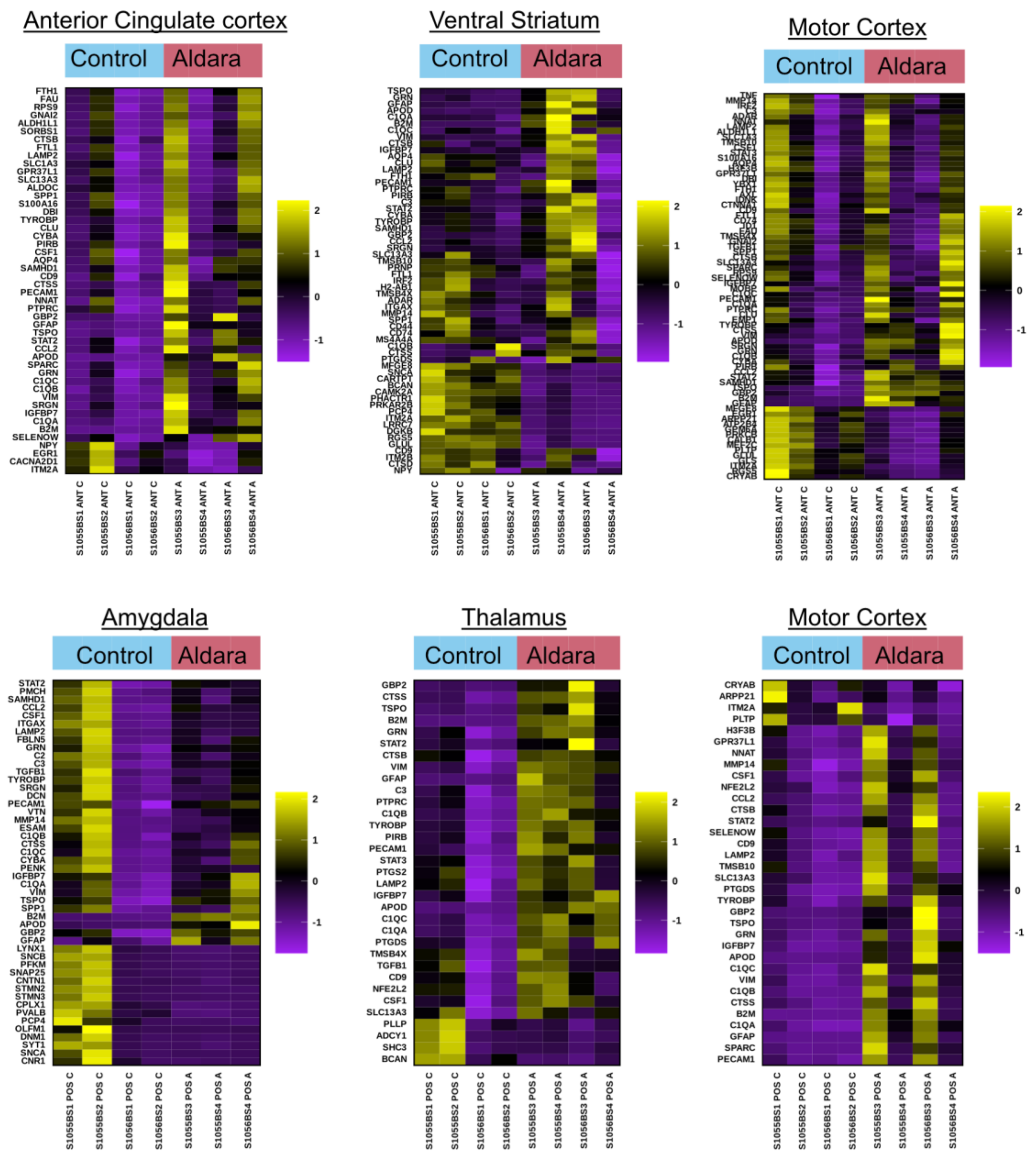

**Supplementary Figure 4: Heatmaps for all significantly up- and down-regulated genes in anterior slices (top row: anterior cingulate cortex, ventral striatum and motor cortex) and posterior slices (bottom row: amygdala, thalamus and motor cortex).** Expression levels shown as raw scaled Z-scores with yellow representing high expression and purple representing low expression, as compared against control samples.

**Supplementary Table 2: All DEGs from ROIs**

| **Anterior cingulate cortex** | | | | | |
|---|---|---|---|---|---|
| Upregulated genes | $Log_2$fold | $p_{adj}$ | Downregulated genes | $Log_2$fold | $p_{adj}$ |
| B2M | 4.16 | 3.09E-46 | EGR1 | -1.03 | 1.14E-06 |
| GFAP | 3.6 | 1.95E-29 | CACNA2D1 | -0.56 | 0.00071441 |
| APOD | 3.28 | 7.56E-27 | NPY | -0.58 | 0.00057033 |
| TSPO | 2.54 | 8.68E-12 | ITM2A | -1.1 | 5.84E-07 |
| C1QA | 1.98 | 9.15E-20 | | | |
| VIM | 1.77 | 2.27E-17 | | | |

| | | | | | |
|---|---|---|---|---|---|
| C1QC | 1.55 | 9.15E-20 | | | |
| GRN | 1.49 | 1.07E-13 | | | |
| CTSS | 1.49 | 4.73E-10 | | | |
| GBP2 | 1.39 | 0.00303321 | | | |
| IGFBP7 | 1.36 | 2.69E-10 | | | |
| PECAM1 | 1.26 | 3.33E-05 | | | |
| C1QB | 1.24 | 2.43E-08 | | | |
| CYBA | 1.21 | 8.62E-06 | | | |
| STAT2 | 1.14 | 1.69E-05 | | | |
| SPARC | 1.13 | 2.44E-08 | | | |
| TYROBP | 1.1 | 2.43E-08 | | | |
| SRGN | 1.04 | 0.00013423 | | | |
| NNAT | 0.98 | 3.33E-05 | | | |
| SAMHD1 | 0.98 | 0.00057033 | | | |
| CCL2 | 0.96 | 0.00760267 | | | |
| CLU | 0.87 | 1.61E-05 | | | |
| SLC13A3 | 0.83 | 1.69E-06 | | | |
| DBI | 0.82 | 7.80E-07 | | | |
| CSF1 | 0.77 | 0.00021127 | | | |
| SPP1 | 0.73 | 0.00159379 | | | |
| LAMP2 | 0.73 | 5.99E-05 | | | |
| RPS9 | 0.71 | 9.86E-07 | | | |
| FTH1 | 0.69 | 4.68E-05 | | | |
| SELENOW | 0.68 | 0.00195184 | | | |
| CTSB | 0.66 | 0.00070396 | | | |
| S100A16 | 0.65 | 0.00037275 | | | |
| FTL1 | 0.65 | 9.23E-05 | | | |
| ALDOC | 0.63 | 2.89E-05 | | | |
| SORBS1 | 0.63 | 0.00037275 | | | |
| CD9 | 0.61 | 0.00475996 | | | |
| SLC1A3 | 0.6 | 0.0004245 | | | |
| PIRB | 0.57 | 0.01536216 | | | |
| GNAI2 | 0.56 | 0.00016093 | | | |
| ALDH1L1 | 0.53 | 0.00345656 | | | |
| GPR37L1 | 0.52 | 0.00038407 | | | |
| AQP4 | 0.52 | 0.00892367 | | | |
| FAU | 0.52 | 0.00085985 | | | |
| PTPRC | 0.51 | 0.03278761 | | | |
| **Ventral striatum** | | | | | |
| Upregulated genes | $Log_2 fold$ | $p_{adj}$ | Downregulated genes | $Log_2 fold$ | $p_{adj}$ |
| B2M | 4.73 | 1.97E-61 | GLUL | -0.57 | 0.00460027 |
| GFAP | 3.17 | 2.01E-25 | RGS5 | -0.64 | 0.0007491 |
| APOD | 2.99 | 4.43E-33 | DGKB | -0.6 | 0.0015271 |
| TSPO | 2.48 | 3.75E-14 | PRKAR2B | -0.55 | 0.00221532 |
| GBP2 | 2.47 | 4.88E-10 | CARTPT | -1.06 | 0.00171025 |
| GRN | 1.95 | 1.86E-17 | PHACTR1 | -0.67 | 0.00010895 |
| C1QA | 1.89 | 4.47E-24 | PCP4 | -0.76 | 0.00012167 |
| VIM | 1.85 | 9.83E-09 | CAMK2A | -0.54 | 0.00455609 |
| C1QC | 1.62 | 6.46E-11 | BCAN | -0.62 | 0.0002814 |
| STAT2 | 1.56 | 6.35E-06 | LRRC7 | -0.58 | 0.00106488 |
| PECAM1 | 1.55 | 2.27E-08 | SNCA | -1.2 | 1.34E-06 |
| CTSB | 1.47 | 3.04E-11 | MFGE8 | -0.64 | 0.00057496 |
| CYBA | 1.47 | 2.49E-07 | ITM2A | -1.04 | 3.49E-09 |

| | | | | | |
|---|---|---|---|---|---|
| SRGN | 1.46 | 2.42E-05 | | | |
| TYROBP | 1.44 | 6.85E-09 | | | |
| IGFBP7 | 1.38 | 2.63E-08 | | | |
| SAMHD1 | 1.36 | 0.0003277 | | | |
| CCL2 | 1.29 | 0.00485174 | | | |
| PTPRC | 1.22 | 8.94E-05 | | | |
| C3 | 1.21 | 0.0003277 | | | |
| SLC13A3 | 1.03 | 5.14E-07 | | | |
| PIRB | 1.01 | 0.00085827 | | | |
| LAMP2 | 0.98 | 1.89E-05 | | | |
| PTGDS | 0.94 | 0.00312518 | | | |
| CD74 | 0.85 | 0.00053667 | | | |
| ITGAX | 0.85 | 0.00057496 | | | |
| TMSB4X | 0.82 | 2.51E-14 | | | |
| FTH1 | 0.78 | 0.00018619 | | | |
| NPY | 0.78 | 0.00125181 | | | |
| PRNP | 0.71 | 6.18E-06 | | | |
| SPP1 | 0.69 | 0.0026455 | | | |
| CLU | 0.68 | 0.00078807 | | | |
| H2-AB1 | 0.63 | 0.00327031 | | | |
| CTSS | 0.63 | 0.01896493 | | | |
| C1QB | 0.61 | 0.01192619 | | | |
| CTSD | 0.6 | 0.0003277 | | | |
| AQP4 | 0.58 | 0.00972004 | | | |
| ADAR | 0.58 | 0.00455609 | | | |
| CD9 | 0.58 | 0.00171025 | | | |
| ITM2B | 0.57 | 3.91E-05 | | | |
| MS4A4A | 0.57 | 0.00558098 | | | |
| TMSB10 | 0.57 | 0.0103642 | | | |
| FTL1 | 0.57 | 0.00171025 | | | |
| CD44 | 0.56 | 0.00787497 | | | |
| MMP14 | 0.52 | 0.01693153 | | | |
| IRF2 | 0.51 | 0.01419563 | | | |
| **Motor cortex (anterior)** | | | | | |
| Upregulated genes | $Log_2$fold | $p_{adj}$ | Downregulated genes | $Log_2$fold | $p_{adj}$ |
| B2M | 4.04 | 6.95E-58 | ATP2B4 | -0.54 | 0.00069387 |
| GFAP | 3.59 | 2.93E-17 | GLS | -0.64 | 7.82E-05 |
| APOD | 3.47 | 9.03E-68 | GLUL | -0.58 | 1.00E-05 |
| TSPO | 2.67 | 1.68E-25 | RGS5 | -0.69 | 5.15E-05 |
| GBP2 | 2.35 | 1.25E-06 | MEF2C | -0.55 | 0.00210419 |
| VIM | 1.96 | 9.85E-30 | EGR1 | -0.55 | 0.00546754 |
| C1QA | 1.78 | 1.47E-33 | PLTP | -0.53 | 0.0046691 |
| CTSS | 1.65 | 3.78E-16 | CALB1 | -0.52 | 0.00539961 |
| GRN | 1.59 | 1.00E-18 | MFGE8 | -0.98 | 2.94E-09 |
| PECAM1 | 1.49 | 5.81E-11 | PRKCB | -0.55 | 6.91E-06 |
| CCL2 | 1.47 | 8.49E-09 | GPM6A | -0.57 | 6.91E-06 |
| STAT2 | 1.42 | 3.19E-08 | ARPP21 | -0.65 | 0.00012424 |
| C1QC | 1.41 | 3.77E-25 | CRYAB | -0.67 | 5.52E-06 |
| TYROBP | 1.4 | 5.64E-12 | ITM2A | -1.16 | 8.41E-11 |
| IGFBP7 | 1.36 | 9.79E-23 | | | |
| SAMHD1 | 1.3 | 1.18E-06 | | | |
| CYBA | 1.29 | 1.49E-08 | | | |
| SRGN | 1.24 | 1.30E-09 | | | |

| | | | | | |
|---|---|---|---|---|---|
| PTPRC | 1.17 | 1.02E-05 | | | |
| C1QB | 1.13 | 4.59E-12 | | | |
| SLC13A3 | 1.05 | 3.28E-09 | | | |
| CLU | 1.03 | 2.42E-09 | | | |
| PIRB | 0.95 | 8.85E-05 | | | |
| SPARC | 0.95 | 2.61E-10 | | | |
| NNAT | 0.95 | 1.88E-07 | | | |
| CTSB | 0.92 | 1.16E-14 | | | |
| LAMP2 | 0.89 | 2.42E-09 | | | |
| RPS9 | 0.87 | 1.03E-14 | | | |
| SELENOW | 0.84 | 1.25E-06 | | | |
| CD9 | 0.83 | 1.18E-06 | | | |
| ADAR | 0.82 | 7.28E-07 | | | |
| STAT3 | 0.79 | 3.95E-06 | | | |
| SPP1 | 0.76 | 8.55E-06 | | | |
| SLC1A3 | 0.75 | 1.16E-06 | | | |
| C3 | 0.74 | 0.00084481 | | | |
| ALDH1L1 | 0.74 | 3.45E-05 | | | |
| CSF1 | 0.74 | 1.18E-05 | | | |
| TNF | 0.71 | 0.00093044 | | | |
| EMP1 | 0.71 | 8.96E-05 | | | |
| FTH1 | 0.7 | 4.61E-07 | | | |
| S100A16 | 0.7 | 3.95E-06 | | | |
| ID1 | 0.68 | 2.66E-08 | | | |
| TMSB10 | 0.67 | 1.00E-05 | | | |
| H3F3B | 0.66 | 1.95E-05 | | | |
| FTL1 | 0.63 | 1.02E-05 | | | |
| FAU | 0.62 | 1.04E-07 | | | |
| AXL | 0.61 | 0.00135034 | | | |
| IDNK | 0.61 | 0.00057177 | | | |
| AQP4 | 0.61 | 6.38E-07 | | | |
| MOBP | 0.6 | 0.00024401 | | | |
| DBI | 0.6 | 0.00248991 | | | |
| GNAI2 | 0.58 | 1.02E-05 | | | |
| YBX1 | 0.57 | 2.17E-05 | | | |
| IRF2 | 0.55 | 3.72E-05 | | | |
| TMSB4X | 0.54 | 0.00601043 | | | |
| TGFB1 | 0.54 | 1.90E-06 | | | |
| CTNNA1 | 0.52 | 0.00124062 | | | |
| MMP14 | 0.52 | 0.00109259 | | | |
| CD74 | 0.51 | 0.01301814 | | | |
| **Amygdala** | | | | | |
| Upregulated genes | $Log_2$fold | $p_{adj}$ | Downregulated genes | $Log_2$fold | $p_{adj}$ |
| DCN | 0.74 | 4.23E-05 | SYT1 | -0.53 | 0.00612964 |
| PMCH | 0.6 | 0.00340748 | SNCB | -0.83 | 6.70E-06 |
| SRGN | 0.84 | 0.00027299 | CNTN1 | -0.57 | 0.00167533 |
| STAT2 | 1.29 | 2.16E-09 | LYNX1 | -0.52 | 0.00167533 |
| CCL2 | 1.11 | 3.42E-08 | PFKM | -0.84 | 1.27E-06 |
| GFAP | 1.64 | 5.20E-05 | PVALB | -0.58 | 0.0166623 |
| GRN | 1.41 | 1.91E-18 | PCP4 | -0.6 | 0.02278964 |
| PECAM1 | 1.47 | 2.00E-09 | DNM1 | -0.53 | 0.00502125 |
| VTN | 0.57 | 0.0166623 | OLFM1 | -0.82 | 0.00767103 |
| FBLN5 | 0.9 | 7.68E-06 | SNAP25 | -0.81 | 1.92E-05 |

| | | | | | |
|---|---|---|---|---|---|
| MMP14 | 0.57 | 0.00311835 | STMN3 | -0.56 | 0.00311835 |
| TSPO | 1.93 | 5.28E-23 | STMN2 | -0.6 | 0.000397 |
| APOD | 0.97 | 0.01071322 | CNR1 | -0.68 | 0.00844085 |
| C2 | 0.74 | 0.0001729 | CPLX1 | -0.95 | 7.81E-05 |
| C3 | 1.27 | 1.91E-18 | SNCA | -0.52 | 0.03788967 |
| B2M | 3.96 | 1.72E-37 | | | |
| SAMHD1 | 0.68 | 0.00222605 | | | |
| VIM | 1.71 | 1.75E-12 | | | |
| CSF1 | 0.64 | 0.00222605 | | | |
| CTSS | 0.69 | 0.00925221 | | | |
| GBP2 | 1.96 | 1.91E-18 | | | |
| C1QA | 1.75 | 5.17E-13 | | | |
| C1QB | 0.89 | 0.00099376 | | | |
| C1QC | 1.38 | 6.48E-10 | | | |
| PENK | 0.84 | 0.00783589 | | | |
| IGFBP7 | 1.44 | 4.73E-07 | | | |
| SPP1 | 1.16 | 1.28E-05 | | | |
| ITGAX | 0.83 | 3.86E-06 | | | |
| TGFB1 | 0.65 | 0.00059633 | | | |
| TYROBP | 0.99 | 2.16E-09 | | | |
| CYBA | 0.85 | 0.00111294 | | | |
| ESAM | 0.58 | 0.00434284 | | | |
| LAMP2 | 0.51 | 0.00721704 | | | |
| **Motor cortex** | | | | | |
| Upregulated genes | $Log_2$fold | $p_{adj}$ | Downregulated genes | $Log_2$fold | $p_{adj}$ |
| GPR37L1 | 0.57 | 0.01175678 | PLTP | -0.55 | 0.03171689 |
| STAT2 | 0.81 | 0.00655707 | ARPP21 | -0.57 | 0.00365697 |
| CCL2 | 0.88 | 0.00072112 | CRYAB | -0.8 | 0.0010029 |
| GFAP | 3.88 | 7.11E-54 | ITM2A | -0.69 | 0.01745563 |
| GRN | 1.2 | 9.92E-06 | | | |
| H3F3B | 0.57 | 0.00274015 | | | |
| PECAM1 | 1.24 | 3.77E-05 | | | |
| SPARC | 0.65 | 0.01302212 | | | |
| CTSB | 0.67 | 0.00274015 | | | |
| MMP14 | 0.56 | 0.0197414 | | | |
| TSPO | 2.43 | 1.83E-08 | | | |
| APOD | 3.14 | 1.16E-18 | | | |
| B2M | 4.3 | 2.36E-56 | | | |
| NFE2L2 | 0.8 | 0.00274015 | | | |
| NNAT | 0.98 | 7.57E-08 | | | |
| PTGDS | 0.78 | 0.00274015 | | | |
| SLC13A3 | 0.88 | 0.00063577 | | | |
| VIM | 1.02 | 0.00027725 | | | |
| CSF1 | 0.81 | 0.00120574 | | | |
| CTSS | 0.95 | 0.00016822 | | | |
| GBP2 | 2.04 | 0.00108289 | | | |
| C1QA | 1.96 | 6.77E-16 | | | |
| C1QB | 0.74 | 0.00556071 | | | |
| C1QC | 1.07 | 0.00013367 | | | |
| IGFBP7 | 0.76 | 0.00933244 | | | |
| CD9 | 0.55 | 0.00725131 | | | |
| TMSB10 | 0.53 | 0.01493738 | | | |
| SELENOW | 0.75 | 0.00043912 | | | |

| | | | | | |
|---|---|---|---|---|---|
| TYROBP | 0.74 | 0.00348473 | | | |
| LAMP2 | 0.52 | 0.011999 | | | |
| **Thalamus** | | | | | |
| Upregulated genes | $Log_2$fold | $p_{adj}$ | Downregulated genes | $Log_2$fold | $p_{adj}$ |
| PTGS2 | 0.66 | 0.00011581 | ADCY1 | -0.92 | 1.75E-07 |
| PTPRC | 1.32 | 3.23E-14 | SHC3 | -0.52 | 0.00140034 |
| STAT2 | 0.59 | 0.03223967 | BCAN | -0.88 | 0.00055621 |
| GFAP | 2.98 | 8.02E-22 | PLLP | -0.6 | 0.00099902 |
| GRN | 1.21 | 2.41E-12 | | | |
| PECAM1 | 1.4 | 3.86E-16 | | | |
| STAT3 | 0.68 | 0.00092549 | | | |
| CTSB | 0.98 | 3.01E-10 | | | |
| TSPO | 2.12 | 2.84E-06 | | | |
| APOD | 2.62 | 1.21E-19 | | | |
| C3 | 1.08 | 8.90E-10 | | | |
| B2M | 4.39 | 3.33E-55 | | | |
| NFE2L2 | 0.56 | 0.00048558 | | | |
| PTGDS | 0.7 | 0.00475963 | | | |
| SLC13A3 | 0.51 | 0.00048071 | | | |
| VIM | 1.48 | 6.89E-16 | | | |
| CSF1 | 0.72 | 0.00048071 | | | |
| CTSS | 1.38 | 1.86E-07 | | | |
| GBP2 | 2 | 0.00015461 | | | |
| C1QA | 1.76 | 3.31E-23 | | | |
| C1QB | 1.12 | 1.86E-07 | | | |
| C1QC | 1.45 | 4.36E-13 | | | |
| IGFBP7 | 1.05 | 0.00011196 | | | |
| CD9 | 0.65 | 0.00079049 | | | |
| PIRB | 0.8 | 0.00055621 | | | |
| TGFB1 | 0.56 | 3.21E-08 | | | |
| TYROBP | 1.04 | 3.06E-07 | | | |
| LAMP2 | 0.55 | 0.00135966 | | | |
| TMSB4X | 0.59 | 0.0001161 | | | |

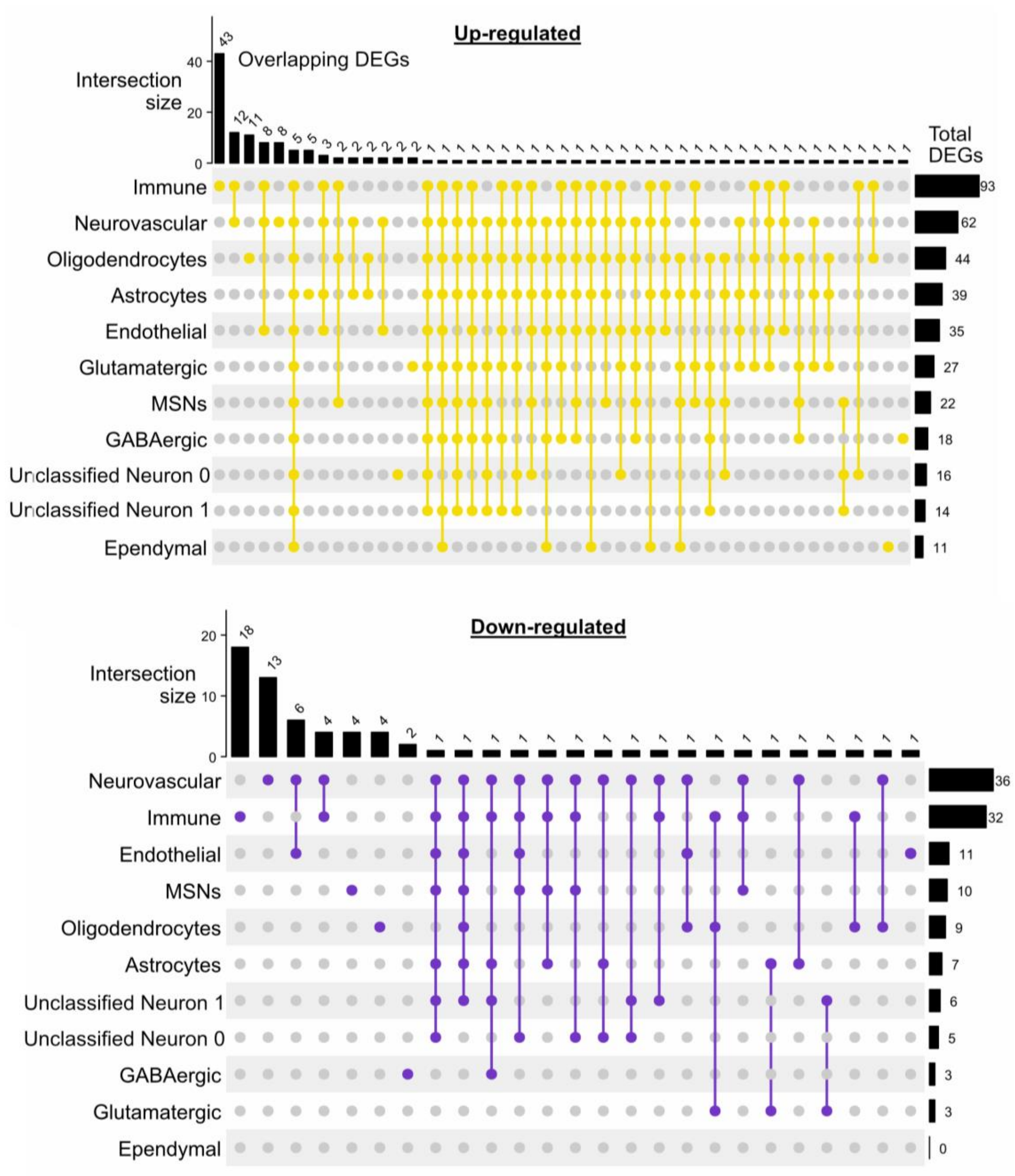

**Supplementary Figure 5: Upset plots for pseudo-bulk analysis of anterior sections. Upregulated (top) and downregulated (bottom) genes and their expression in different cell types.** Top axis shown number of overlapping DEGs in different cell types (single dot = unique DEG, linked dots = DEG present in the highlighted cell type); right axis shows total number of DEG per each cell type.

**Supplementary Table 3: combination summary of up- and down-regulated genes in anterior structures**

| **Combination summary upregulated genes (anterior sections)** | |
|---|---|
| **Cell Types** | **Genes** |
| Astrocytes, Unclassified Neuron 0, Unclassified Neuron 1, Endothelial, Ependymal, GABAergic, Glutamatergic, Immune, MSNs, Neurovascular, Oligodendrocytes | Apod, B2m, Gfap, Stat2, Vim |

| | |
|---|---|
| Astrocytes, Unclassified Neuron 1, Endothelial, Ependymal, GABAergic, Glutamatergic, Immune, MSNs, Neurovascular, Oligodendrocytes | Gbp2 |
| Astrocytes, Unclassified Neuron 0, Unclassified Neuron 1, Endothelial, GABAergic, Glutamatergic, Immune, MSNs, Neurovascular, Oligodendrocytes | Grn |
| Astrocytes, Unclassified Neuron 1, Endothelial, GABAergic, Glutamatergic, Immune, MSNs, Neurovascular, Oligodendrocytes | Tspo |
| Astrocytes, Unclassified Neuron 0, Unclassified Neuron 1, GABAergic, Glutamatergic, Immune, MSNs, Neurovascular, Oligodendrocytes | C1qa |
| Astrocytes, Unclassified Neuron 1, Endothelial, GABAergic, Glutamatergic, Immune, Neurovascular, Oligodendrocytes | Selenow |
| Astrocytes, Unclassified Neuron 0, Unclassified Neuron 1, GABAergic, Glutamatergic, MSNs, Neurovascular, Oligodendrocytes | C1qc |
| Astrocytes, Endothelial, GABAergic, Immune, MSNs, Neurovascular, Oligodendrocytes | Lamp2 |
| Astrocytes, Endothelial, GABAergic, Glutamatergic, Immune, Neurovascular, Oligodendrocytes | Samhd1 |
| Astrocytes, Endothelial, Ependymal, GABAergic, Glutamatergic, Neurovascular, Oligodendrocytes | Sparc |
| Astrocytes, Unclassified Neuron 0, Endothelial, Immune, MSNs, Neurovascular, Oligodendrocytes | Ctsb |
| Astrocytes, Unclassified Neuron 0, Unclassified Neuron 1, Glutamatergic, Immune, Neurovascular, Oligodendrocytes | Adar |
| Endothelial, GABAergic, Glutamatergic, MSNs, Neurovascular, Oligodendrocytes | C1qb |
| Unclassified Neuron 0, Endothelial, Glutamatergic, Immune, Neurovascular, Oligodendrocytes | Tmsb10 |
| Astrocytes, Endothelial, Immune, MSNs, Neurovascular, Oligodendrocytes | Igfbp7 |

| | |
|---|---|
| Astrocytes, Endothelial, Ependymal, Immune, Neurovascular, Oligodendrocytes | Fth1 |
| cluster1, GABAergic, Glutamatergic, MSNs, Oligodendrocytes | Idnk |
| Astrocytes, Glutamatergic, Immune, MSNs, Neurovascular | Ctss |
| Astrocytes, Ependymal, Glutamatergic, MSNs, Oligodendrocytes | Clu |
| Astrocytes, Endothelial, Immune, Neurovascular, Oligodendrocytes | Srgn |
| Astrocytes, Endothelial, Ependymal, Immune, Neurovascular | C3 |
| GABAergic, Glutamatergic, MSNs, Oligodendrocytes | Ptgds |
| Endothelial, Immune, Neurovascular, Oligodendrocytes | Rps9 |
| Endothelial, Glutamatergic, Immune, Neurovascular | Pecam1 |
| Astrocytes, Glutamatergic, Immune, Oligodendrocytes | Stat3 |
| Astrocytes, Endothelial, Immune, Neurovascular | H3f3b, Hspa8, Tmsb4x |
| Astrocytes, Endothelial, Glutamatergic, Neurovascular | Cd9 |
| Astrocytes, Unclassified Neuron 0, MSNs, Oligodendrocytes | Itm2b |
| Immune, MSNs, Oligodendrocytes | Irf2, Slc13a3 |
| Endothelial, Immune, Neurovascular | Ccl2, Emp1, Fau, Ftl1, Hsp90ab1, Lyz1/2, Spp1, Ybx1 |
| Unclassified Neuron 0, Unclassified Neuron 1, MSNs | Prnp |
| Astrocytes, Glutamatergic, Oligodendrocytes | Slc1a3 |
| Astrocytes, Glutamatergic, Neurovascular | S100a16 |
| Immune, Oligodendrocytes | Fos |
| Immune, Neurovascular | Arhgef7, Atf6, Cast, Cd44, Cyba, Mmp14, Nfe2l2, Pirb, Psma1, Ptpn1, Ptprc, Tyrobp |
| Endothelial, Neurovascular | Ctsd, Rtn4 |
| Unclassified Neuron 0, Immune | Psap |
| Astrocytes, Oligodendrocytes | Aldh1l1, Fabp7 |
| Astrocytes, Neurovascular | Csf1, Gpnmb |
| Oligodendrocytes | Adora2a, Bdnf, Brwd1, Cdc14b, Grik2, Lpar6, Mt3, Nap1l5, Nlgn4l, Slc2a13, Tnrc6a |
| Neurovascular | Adam10, Cd74, Cyfip1, Dcn, Malat1, Pros1, Ptgs2, Rapgef1 |
| Immune | Acer3, C2, C5ar1, Cldn5, Cpa3, Crip1, Dync1h1, Fn1, Fnip2, Fpr2, Fus, Gnai2, Grb2, Id1, Id2, Il10, Itgax, Jak2, Jun, Ldha, Lilrb4a/b, Man2a1, Ms4a4a, Msr1, Myl6, P2rx4, Pgd, |

| | |
|---|---|
| | Rab1a, Rab31, Ranbp2, Rapgef2, Rock1, Sem1, Slc11a2, Slc2a1, Sos1, Taldo1, Tkt, Tnf, Tubb5, Vcan, Wwp2, Xpo1 |
| Glutamatergic | Mobp, Ndrg2 |
| GABAergic | Dbi |
| Ependymal | App |
| Unclassified Neuron 0 | Atp2a2, Gria2 |
| Astrocytes | Aqp4, Clec7a, Gabrb2, H2-Ab1, Sorbs1 |
| **Combination summary downregulated genes (anterior sections)** | |
| **Cell Types** | **Genes** |
| Astrocytes, Unclassified Neuron 1, Endothelial, Immune, MSNs, Neurovascular, Oligodendrocytes | Rgs5 |
| Astrocytes, Unclassified Neuron 0, Unclassified Neuron 1, Endothelial, Immune, MSNs, Neurovascular | Glul |
| Unclassified Neuron 0, Endothelial, Immune, MSNs, Neurovascular | Slc1a2 |
| Astrocytes, Unclassified Neuron 1, GABAergic, Immune, Neurovascular | Adcy1 |
| Unclassified Neuron 0, Immune, MSNs, Neurovascular | Pcp4 |
| Astrocytes, Immune, MSNs, Neurovascular | Bcan |
| Immune, MSNs, Neurovascular | Snca |
| Glutamatergic, Immune, Oligodendrocytes | Csf1r |
| Endothelial, Neurovascular, Oligodendrocytes | Ndrg1 |
| Unclassified Neuron 1, Immune, Neurovascular | Grm3 |
| Unclassified Neuron 0, Unclassified Neuron 1, Neurovascular | Scd2 |
| Astrocytes, Unclassified Neuron 0, Neurovascular | Gng7 |
| Neurovascular, Oligodendrocytes | Pllp |
| Immune, Oligodendrocytes | Plp1 |
| Immune, Neurovascular | Ctxn1, Mbp, Nrgn, Ptgds |
| Endothelial, Neurovascular | Edn3, Igf1r, Itm2a, Pltp, Slc39a10, Slc7a5 |
| Unclassified Neuron 1, Glutamatergic | Gls |
| Astrocytes, Neurovascular | Phactr1 |
| Astrocytes, Glutamatergic | Mfge8 |
| Oligodendrocytes | Clasp2, Cryab, Gnai1, Phlpp1 |
| Neurovascular | Atp1a2, Camk2a, Camk4, Chrm1, Gpm6a, Gpm6b, Myl9, Pomc, Ppp2r2c, Prkag2, Prkcb, Snap25, Tspan7 |
| MSNs | Cartpt, Foxo1, Gap43, Plpp3 |

| | |
|---|---|
| Immune | Adgrg1, Arpp21, Atg2b, Camk2b, Cfh, Cst3, Cx3cr1, Hexb, Mef2c, Mertk, P2ry12, Prkca, Sall1, Selplg, Snn, Tgfbr1, Tmem119, Trem2 |
| GABAergic | Cxcl14, Egr1 |
| Endothelial | Ntrk2 |

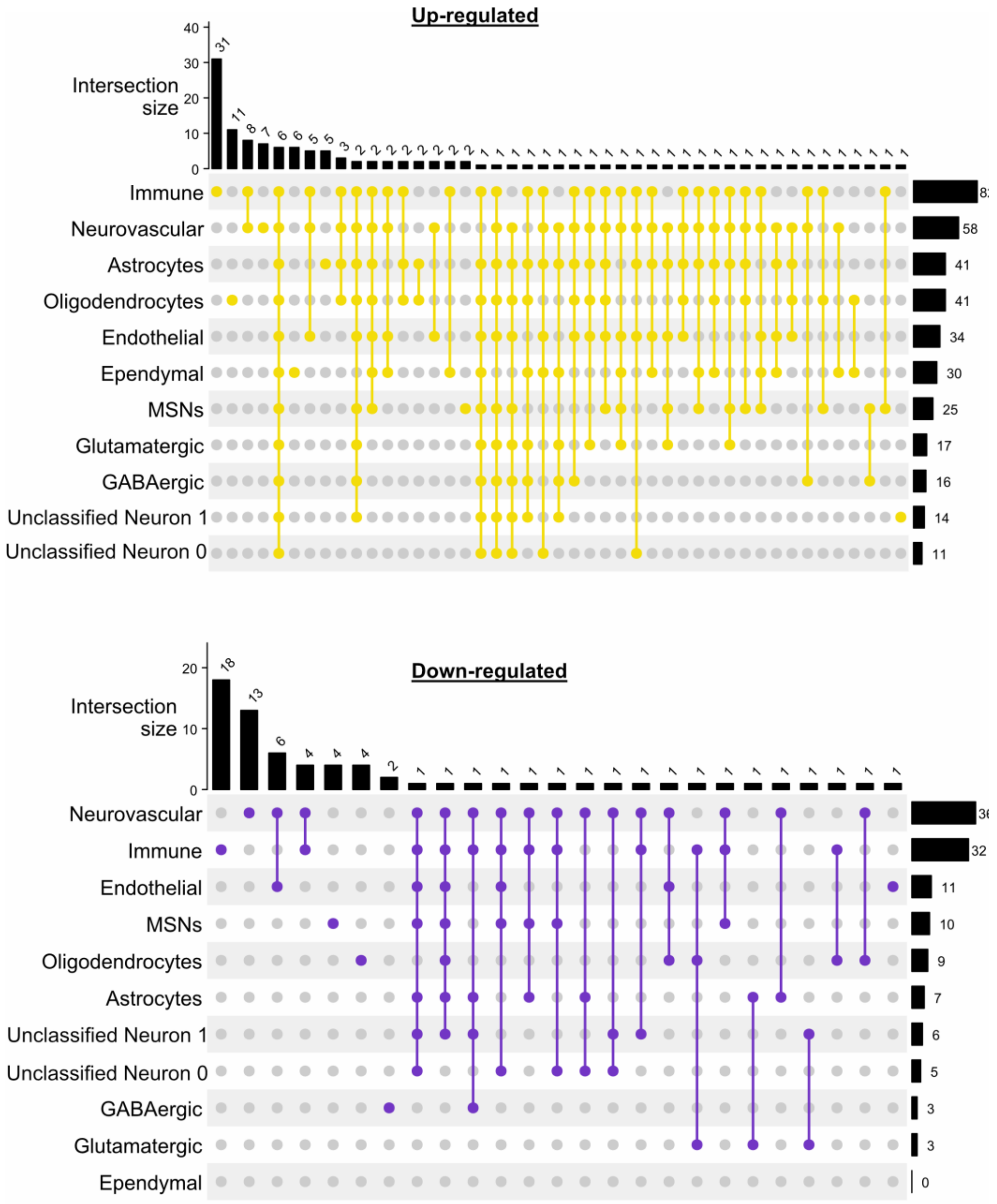

**Supplementary Figure 6: Upset plots for pseudo bulk analysis of posterior sections.** Upregulated (top) and downregulated (bottom) genes and their expression in different cell types. Top axis shown number of overlapping DEGs in different cell types (single dot = unique DEG, linked dots = DEG present in the highlighted cell type); right axis shows total number of DEG per each cell type.

**Supplementary Table 4: combination summary of up- and down-regulated genes in posterior structures.**

| CellTypes | Genes |
|---|---|

| | |
|---|---|
| Astrocytes, Unclassified Neuron 0, Unclassified Neuron 1, Endothelial, Ependymal, GABAergic, Glutamatergic, Immune, MSNs, Neurovascular, Oligodendrocytes | Apod, B2m, Gfap, Grn, Tspo, Vim |
| Astrocytes, Unclassified Neuron 0, Unclassified Neuron 1, Endothelial, GABAergic, Glutamatergic, Immune, MSNs, Neurovascular, Oligodendrocytes | C1qa |
| Astrocytes, Unclassified Neuron 0, Unclassified Neuron 1, Endothelial, Ependymal, GABAergic, Glutamatergic, Immune, MSNs, Oligodendrocytes | Itm2b |
| Astrocytes, cluster1, Endothelial, GABAergic, Glutamatergic, Immune, MSNs, Neurovascular, Oligodendrocytes | Gbp2, Stat2 |
| Astrocytes, Unclassified Neuron 0, Unclassified Neuron 1, Endothelial, GABAergic, Glutamatergic, MSNs, Neurovascular, Oligodendrocytes | C1qc |
| Astrocytes, Endothelial, Ependymal, Immune, MSNs, Neurovascular, Oligodendrocytes | C3, Ctsb |
| Astrocytes, Unclassified Neuron 1, Ependymal, GABAergic, Glutamatergic, Immune, Oligodendrocytes | Selenow |
| Endothelial, Ependymal, Glutamatergic, Immune, MSNs, Neurovascular | Tmsb10 |
| Astrocytes, Endothelial, Immune, MSNs, Neurovascular, Oligodendrocytes | Lamp2 |
| Astrocytes, Endothelial, Glutamatergic, Immune, Neurovascular, Oligodendrocytes | Pecam1 |
| Astrocytes, Endothelial, GABAergic, Immune, Neurovascular, Oligodendrocytes | Tyrobp |
| Astrocytes, Unclassified Neuron 1, Ependymal, GABAergic, Glutamatergic, Neurovascular | Csf1 |
| Astrocytes, Unclassified Neuron 0, Endothelial, Ependymal, Immune, Neurovascular | H3f3b |
| Endothelial, Ependymal, Immune, MSNs, Neurovascular | Ccl2 |
| Astrocytes, Immune, MSNs, Neurovascular, Oligodendrocytes | C1qb |
| Astrocytes, Glutamatergic, Immune, MSNs, Neurovascular | Nfe2l2 |

| | |
|---|---|
| Astrocytes, Ependymal, Immune, Neurovascular, Oligodendrocytes | Adar |
| Astrocytes, Ependymal, Immune, MSNs, Neurovascular | Ctss |
| Astrocytes, Endothelial, Immune, Neurovascular, Oligodendrocytes | Igfbp7 |
| Astrocytes, Endothelial, Glutamatergic, MSNs, Neurovascular | Cd9 |
| Astrocytes, Endothelial, Ependymal, Immune, Neurovascular | Tmsb4x |
| Astrocytes, Unclassified Neuron 0, Endothelial, Immune, Neurovascular | Hspa8 |
| Endothelial, Ependymal, Immune, Neurovascular | Ftl1, Spp1 |
| Astrocytes, Immune, Neurovascular, Oligodendrocytes | Ptprc, Samhd1, Stat3 |
| Astrocytes, Endothelial, Neurovascular, Oligodendrocytes | Gpr37l1 |
| Astrocytes, Endothelial, Ependymal, Neurovascular | C2 |
| Immune, MSNs, Oligodendrocytes | Slc13a3 |
| GABAergic, Immune, Neurovascular | Mmp14 |
| Endothelial, Immune, Neurovascular | Fth1, Ldha, Psma1, Srgn, Ybx1 |
| Astrocytes, Immune, Oligodendrocytes | Fos, Gpnmb |
| Immune, Neurovascular | Cast, Cyba, Gnai2, Itgax, Lyz1/2, Pirb, Tnf, Xpo1 |
| Immune, MSNs | Psap |
| GABAergic, MSNs | Idnk |
| Ependymal, Oligodendrocytes | Crh |
| Ependymal, Neurovascular | Sgk1 |
| Ependymal, Immune | Dcn, Esam |
| Endothelial, Neurovascular | Emp1, Rtn4 |
| Astrocytes, Oligodendrocytes | Aldh1l1, Clu |
| Oligodendrocytes | Bdnf, Brwd1, Cdc14b, Drd2, Ghr, Grik2, Lpar6, Mmp9, Mt1, Nlgn4l, Ptgds |
| Neurovascular | Arhgef7, Ctsd, Cyfip1, Dync1i2, Hspa1b, Ptgs2, Tgfb1 |
| MSNs | Chrna4, Prnp |
| Immune | Acer3, Atf6, C5ar1, Cd44, Cldn5, Crip1, Dync1h1, Fau, Fn1, Fpr2, Grb2, Id1, Id2, Lilrb4a/b, Man2a1, Ms4a4a, Msr1, Myl6, Nkg7, P2rx4, Pgd, Ptpn1, Rab1a, Rab31, Ranbp2, Rps9, Sem1, Slc11a2, Slc2a1, Tkt, Tubb5 |
| Ependymal | Ahcyl1, Cd74, Cspg5, Foxj1, Hsp90aa1, Maob |
| Unclassified Neuron 1 | App |
| Astrocytes | Aqp4, Gabrb2, H2-Ab1, Sorbs1, Sox9 |
| **Combination summary downregulated genes (posterior sections)** | |
| **Cell Types** | **Genes** |
| Astrocytes, Unclassified Neuron 1, Endothelial, Immune, MSNs, Neurovascular, Oligodendrocytes | Rgs5 |

| | |
|---|---|
| Astrocytes, Unclassified Neuron 0, Unclassified Neuron 1, Endothelial, Immune, MSNs, Neurovascular | Glul |
| Unclassified Neuron 0, Endothelial, Immune, MSNs, Neurovascular | Slc1a2 |
| Astrocytes, Unclassified Neuron 1, GABAergic, Immune, Neurovascular | Adcy1 |
| Unclassified Neuron 0, Immune, MSNs, Neurovascular | Pcp4 |
| Astrocytes, Immune, MSNs, Neurovascular | Bcan |
| Immune, MSNs, Neurovascular | Snca |
| Glutamatergic, Immune, Oligodendrocytes | Csf1r |
| Endothelial, Neurovascular, Oligodendrocytes | Ndrg1 |
| Unclassified Neuron 1, Immune, Neurovascular | Grm3 |
| Unclassified Neuron 0, Unclassified Neuron 1, Neurovascular | Scd2 |
| Astrocytes, Unclassified Neuron 0, Neurovascular | Gng7 |
| Neurovascular, Oligodendrocytes | Pllp |
| Immune, Oligodendrocytes | Plp1 |
| Immune, Neurovascular | Ctxn1, Mbp, Nrgn, Ptgds |
| Endothelial, Neurovascular | Edn3, Igf1r, Itm2a, Pltp, Slc39a10, Slc7a5 |
| Unclassified Neuron 1, Glutamatergic | Gls |
| Astrocytes, Neurovascular | Phactr1 |
| Astrocytes, Glutamatergic | Mfge8 |
| Oligodendrocytes | Clasp2, Cryab, Gnai1, Phlpp1 |
| Neurovascular | Atp1a2, Camk2a, Camk4, Chrm1, Gpm6a, Gpm6b, Myl9, Pomc, Ppp2r2c, Prkag2, Prkcb, Snap25, Tspan7 |
| MSNs | Cartpt, Foxo1, Gap43, Plpp3 |
| Immune | Adgrg1, Arpp21, Atg2b, Camk2b, Cfh, Cst3, Cx3cr1, Hexb, Mef2c, Mertk, P2ry12, Prkca, Sall1, Selplg, Snn, Tgfbr1, Tmem119, Trem2 |
| GABAergic | Cxcl14, Egr1 |
| Endothelial | Ntrk2 |
